# Mimicry drives convergence in structural and light transmission features of transparent wings in Lepidoptera

**DOI:** 10.1101/2020.06.30.180612

**Authors:** Charline Pinna, Maëlle Vilbert, Stephan Borensztajn, Willy Daney de Marcillac, Florence Piron-Prunier, Aaron F. Pomerantz, Nipam Patel, Serge Berthier, Christine Andraud, Doris Gomez, Marianne Elias

## Abstract

Müllerian mimicry is a positive interspecific interaction, whereby co-occurring defended prey species share a common aposematic signal. In Lepidoptera, aposematic species typically harbour conspicuous opaque wing colour patterns with convergent optical properties among co-mimetic species. Surprisingly, some aposematic mimetic species have partially transparent wings, raising the questions of whether optical properties of transparent patches are also convergent, and of how transparency is achieved. Here we conducted a comparative study of wing optics, micro and nanostructures in neotropical mimetic clearwing Lepidoptera, using spectrophotometry and microscopy imaging. We show that transparency, as perceived by predators, is convergent among co-mimics. Underlying micro- and nanostructures are also convergent despite a large structural diversity. We reveal that while transparency is primarily produced by microstructure modifications, nanostructures largely influence light transmission, maybe enabling additional fine-tuning in transmission properties. This study shows that transparency might not only enable camouflage but can also be part of aposematic signals.

## Introduction

Lepidoptera (butterflies and moths) are characterized by large wings typically covered by scales, as testified by the name of the order (after the ancient greek *lepís* - scale and *pterón* – wing). Scales can contain pigments or generate interference colours, thereby producing colour patterns across the entire wing. Wing colour patterns are involved in thermoregulation (Dufour et al., 2018; Heidrich et al., 2018), sexual selection (Kemp, 2007) and anti-predator defences, such as crypsis (Cook et al., 2012; Endler, 1984; Webster et al., 2009), masquerade (Skelhorn et al., 2010; Stoddard, 2012), disruptive coloration, and deflection of predator attacks (Vallin et al., 2011). Another type of anti-predator defence in Lepidoptera involving wing colour pattern is aposematism, where the presence of secondary defences is advertised by the means of bright and contrasted colour patterns. Because of the positive frequency-dependent selection incurred on aposematic signals (Greenwood et al. 1989, Chouteau et al. 2016), aposematic species often engage in Müllerian mimetic interactions, whereby species exposed to the same suite of predators converge on the same colour pattern and form mimicry ‘rings’ (Müller, 1879). Co-mimetic species (species that share a common aposematic colour pattern) are often distantly related, implying that the same colour pattern has evolved independently multiple times. Among such co-mimetic lepidopteran species, several studies using visual modelling have shown that analogous colour patches (i. e., those occupying a similar position in the wing and harbouring similar colour) cannot be discriminated by birds, believed to be the main predators (Bybee et al., 2012; Llaurens et al., 2014; Su et al., 2015; Thurman & Seymoure, 2016). Therefore, mimicry selects for convergent (when a trait in different species evolves towards the same value) or advergent (when a trait of a given species evolves towards the trait value in another species) colourations, as perceived by predators.

Surprisingly, although most aposematic Lepidoptera species harbour brightly coloured patterns, some unpalatable, aposematic species exhibit transparent wing patches (McClure et al., 2019). In those species, wing colour pattern typically consists of a mosaic of brightly coloured and transparent patches. Notably, in tropical America, many mimicry rings comprise such transparent species (Beccaloni, 1997; Elias et al., 2008; Willmott et al., 2017). Mimicry among species harbouring transparent patches raises the question of selection for convergence in optical properties, as perceived by predators, in those transparent patches.

A related question is whether transparency in co-mimetic species is achieved by the means of similar structural changes in wings and scales. Previous studies on a handful of species (most of which are not aposematic) have revealed several, non-mutually exclusive means to achieve transparency, through scale modification or scale shedding, with the effect of reducing the total coverage of the chitin membrane by scales. Scales can fall upon adult emergence (Yoshida et al., 1996); they can have a reduced size (Dushkina et al., 2017; Perez Goodwyn et al., 2009) and even resemble bristle (Binetti et al., 2009; Hernández-Chavarría et al., 2004; Perez Goodwyn et al., 2009; Siddique et al., 2015); they can be either flat on the membrane (Perez Goodwyn et al., 2009) or erected (Dushkina et al., 2017; Perez Goodwyn et al., 2009), which also reduces effective membrane coverage by scales. Reducing scale density could also make wings transparent to some extent (Perez Goodwyn et al., 2009). Although this has not been reported in transparent Lepidoptera, transparent scales, such as those covering coloured scales in the opaque butterfly *Graphium sarpedon* (Stavenga et al., 2010), could also be a means to achieve transparency. In addition to scale modifications, the presence of nanostructures on the surface of the wing membrane may enhance transparency through the reduction of light reflection, by generating a gradient of refractive index between the chitin-made membrane and the air (Binetti et al., 2009; Siddique et al., 2015; Yoshida et al., 1997). Yet, so far, no study has compared the microstructures (scales) and nanostructures present in transparent patches across co-mimetic species. Furthermore, the diversity of structures described above may lead to a large range of transparency efficiencies. Exploring the link between structural features and optical properties can shed light on whether and how different structures might achieve similar degrees of transparency in the context of mimicry.

Here, we investigate the transmission properties of transparent patches among co-mimetic butterflies and moths and the structural bases of transparency on 62 Neotropical transparent species, which belong to seven different lepidopteran families and represent 10 distinct mimicry rings. We characterise wing micro- and nanostructures with digital microscopy and scanning electron microscopy (SEM) imaging and measure transmission properties of transparent patches using spectrophotometry in the range of wavelengths 300-700 nm. We implement comparative analyses that account for phylogenetic relatedness, to (1) examine the putative convergence or advergence (hereafter, convergence, for the sake of simplicity) among co-mimetic species in visual appearance of transparent patches as seen by bird predators, (2) identify and examine the putative convergence of structures involved in transparency in the different co-mimetic species and finally (3) explore the links between structural features and transmission properties of transparent patches.

## Results

### Convergence among co-mimics in visual appearance of transparent patches as seen by bird predators

To assess whether transparent patches of co-mimetic species were under selection for convergence due to mimicry, we tested whether these transparent patches were more similar, as perceived by predators, among co-mimetic species than expected (1) at random, and (2) given the phylogeny. The first test, which assesses whether predators have a similar perception of analogous transparent patches in co-mimetic species, informs on the selection on transparent patches incurred by predators. The second test, which accounts for the phylogenetic relationship between species, informs on the underlying process leading to similarity, and specifically on whether any case of similarity among co-mimics detected in the first test is due to shared ancestry or to evolutionary convergence. We used spectrophotometry to measure specular transmittance of the transparent patches, which is a quantitative measurement of transparency. As birds are supposed to be the main predators of butterflies (Brower, 1984), we applied bird perceptual vision modelling on the resulting spectra to calculate the chromatic and achromatic contrasts (respectively dS and dL) for each pair of species from our dataset. If transparent patches among co-mimetic species are more similar than expected at random or given the phylogeny, contrasts between pairs of co-mimetic species are expected to be smaller than predicted at random and given the phylogeny, respectively. We only compared analogous spots (i.e. occupying a similar position on the forewing) between species. The results presented in Figure 1 show that for three spots out of five and across all mimicry rings the difference in achromatic contrast (dL) between co-mimetic species is significantly smaller than expected both at random and given the phylogenetic distance between species (figure 1B), irrespective of the illuminating light or the visual system considered (see Supplementary table 1a and b for results under the full range of conditions). Differences in chromatic contrasts (dS) between co-mimetic species are significantly smaller than expected at random and given the phylogenetic distance between species only for the most proximal spot on the forewing (figure 1B). These results mean that, on average, predators see transparent patches among co-mimetic species as more similar than among species that belong to different mimicry rings. The fact that these tests remain significant with the phylogenetic correction indicates that such similarity in transparent patches is due to convergent evolution. When looking more precisely at similarity between co-mimetic species for each individual mimicry ring (figure 1C), we show that in eight out of ten mimicry rings achromatic contrasts (dL) between co-mimetic species are smaller than expected at random for at least one spot on the forewing (Supplementary table 2a). After accounting for the phylogeny, this figure drops down to four out of ten mimicry rings (Supplementary table 2a). Regarding chromatic contrasts (dS), six mimicry rings out of ten comprise co-mimetic species exhibiting smaller chromatic contrast than expected at random and three out of them comprise co-mimetic species with smaller chromatic contrast than expected given the phylogeny (figure 1C, Supplementary table 2b). These results suggest that in most cases, the similarity in transparent patches between co-mimetic species is due to convergent evolution but we cannot rule out that for some mimicry rings (notably ‘theudelinda’, ‘hewitsoni’, ‘panthyale’) similarity could be due to shared ancestry. Achromatic aspects (achromatic contrast dL) appear more significant than chromatic aspects (chromatic contrast dS) (Figure 1, Supplementary table 1), suggesting that selection may act more on broadband transmittance (which is related to the degree of transparency) than on colour.

**Figure 1.**
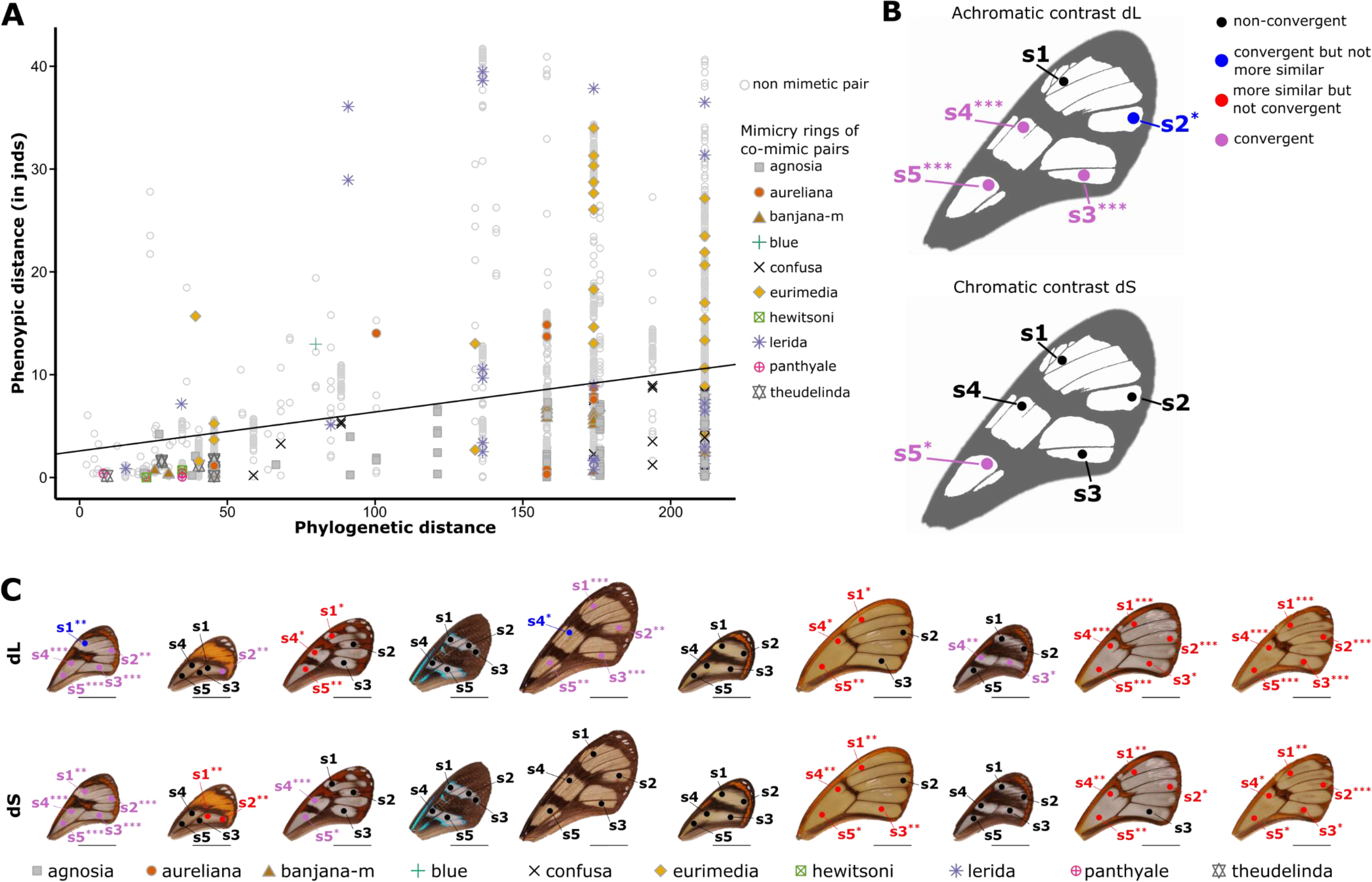
Test of convergence of transmission properties between co-mimetic species. **A. Relation between phenotypic distances (either chromatic or achromatic contrast, expressed in just noticeable differences (jnds), and phylogenetic distances.** Each point represents a pair of species that can be co-mimetic (in this case, the mimicry ring they belong to is represented with a specific symbol, as indicated in the legend) or not (in this case, the point in represented as an open grey circle). The black line represents the predicted value of the linear model linking phenotypic distances with phylogenetic distances. When a point is under the line, the two species represented by that point are more similar that what is expected according to the phylogeny., suggesting that the similarity between these two species is due to convergent evolution. We represent here on this graph the value of achromatic contrast for the 5^th^ spot on the forewing viewed in large gap illuminant by a UVS-type bird but the graphs are similar for all significant tests presented in the Supplementary table 1. **B. Graphical representations of the results of the test of convergence for each spot measured on the forewing, for achromatic (top figure) and chromatic (bottom figure) contrasts, across all mimicry rings.** Each spot measured is localised on the forewing and the results of the test over all mimicry rings are represented by the colour of the spot. Black spots stand for spots that are neither more similar than expected at random nor convergent between co-mimetic species. Blue spots stand for spots that are no more similar than expected at random but which are convergent because they are more similar that what might be expected according to their phylogenetic distance. Red spots stand for spots that are more similar than expected at random but not convergent. Purple spots stand for spots that are both more similar than expected at random and convergent. Tests’ p-values are represented with the following symbols: ‘***’ p < 0.001, ‘**’ p < 0.01, ‘*’ p < 0.05. The same results were obtained irrespective of the bird visual system (either VS or UVS) or the illuminating light (either forest shade or large gap) considered (see Supplementary table 1 for details). **C. Graphical representation of the results of the test of convergent for each spot measured on the forewing, for achromatic (dL, upper row) and chromatic (dS, lower row) contrasts, for each mimicry ring.** The results of the test are represented by the colour of the spot, as indicated above. Tests’ p-values are represented with star symbols as indicated above. The same results were obtained irrespective of the bird visual system (either VS or UVS) or the illuminating light (either forest shade or large gap) considered (see Supplementary table 2 for details).

### Diversity and convergence among co-mimics of structures involved in transparency

Convergence in transmission among co-mimetic species raises the question of the nature and similarity of clearwing microstructures (scales) and nanostructures among co-mimetic species. We therefore explored the diversity of micro- and nanostructures present in the transparent patches in our 62 species. We used digital photonic microscopy and SEM imaging to characterise the structures present in the transparent patches (type, insertion, colour, length, width, and density of scales; type and density of nanostructures; wing membrane thickness).

We found a diversity of microstructural features in transparent patches (Figure 2A). Scales could be coloured (76% of species) or transparent (24%); they could be flat on the membrane (16%) or erected (84%). Scales could be lamellar (i. e., a two-dimension structure, 55% of species), or piliform (45%). In our dataset, piliform scales (mainly bifid) appeared to be almost exclusively found in the Ithomiini tribe, although one erebid species also harboured monofid piliform scales (figure 3).

**Figure 2.**
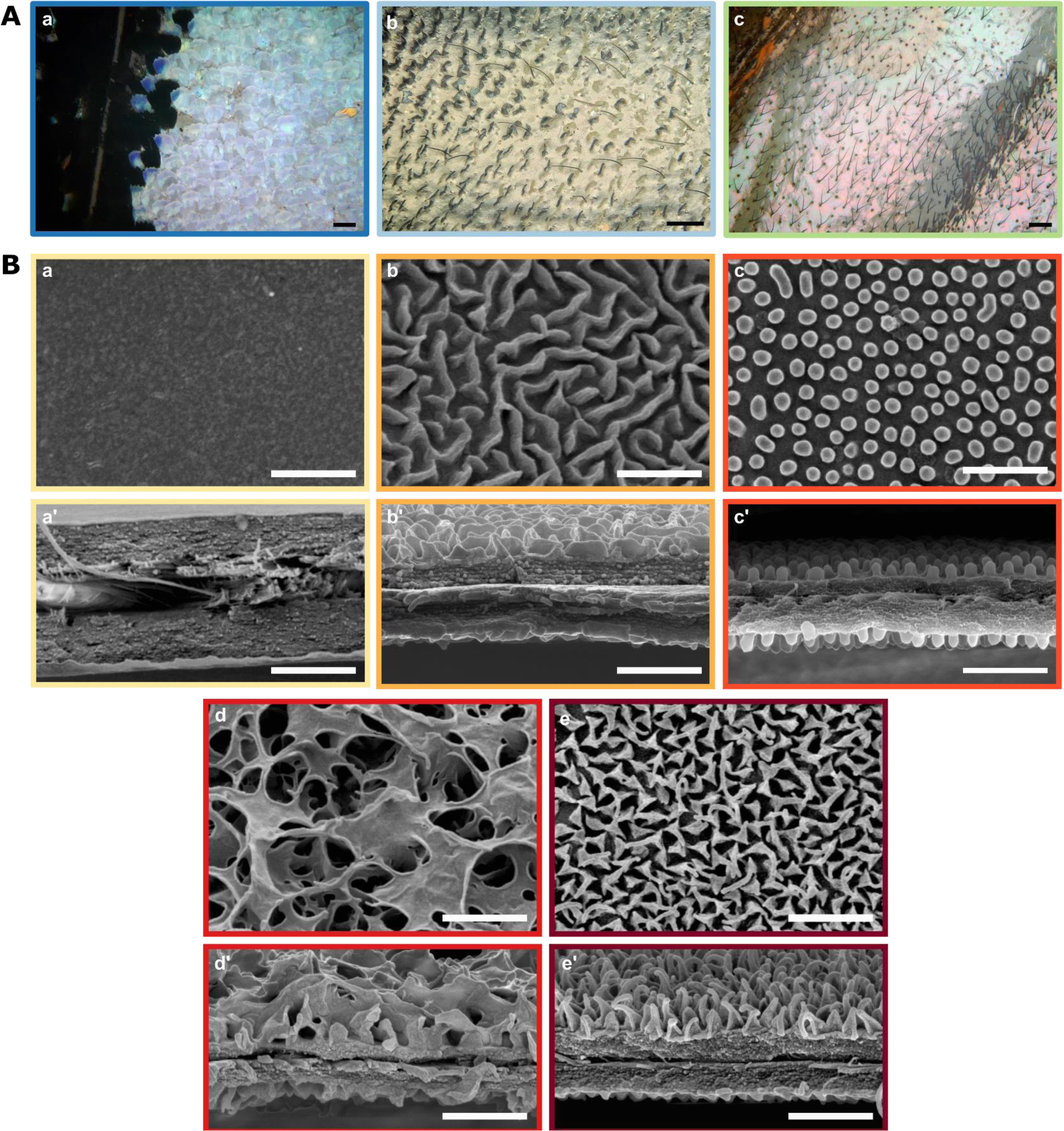
Diversity of micro- and nanostructures involved in transparency. **A. Diversity of microstructures.** a. transparent lamellar scales of *Hypocrita strigifera,* b. erected lamellar scales of *Methona curvifascia* and c. piliform scales of *Hypomenitis ortygia*. Scale bars represent 100 µm. **B. Diversity of nanostructures.** a, b, c, d and e represent top views and a’, b’, c’, d’ and e’ represent cross section of wing membrane. Scale bars represent 1 µm. a, a’. absence of nanostructure in *Methona curvifascia*; b, b’. maze nanostructures of *Megoleria orestilla*; c, c’. nipple nanostructures of *Ithomiola floralis*; d, d’. sponge-like nanostructures of *Oleria onega*; e, e’. pillar nanostructures of *Hypomenitis enigma*. Each coloured frame corresponds to a scale type or nanostructure type, as defined in figure 3.

**Figure 3.**
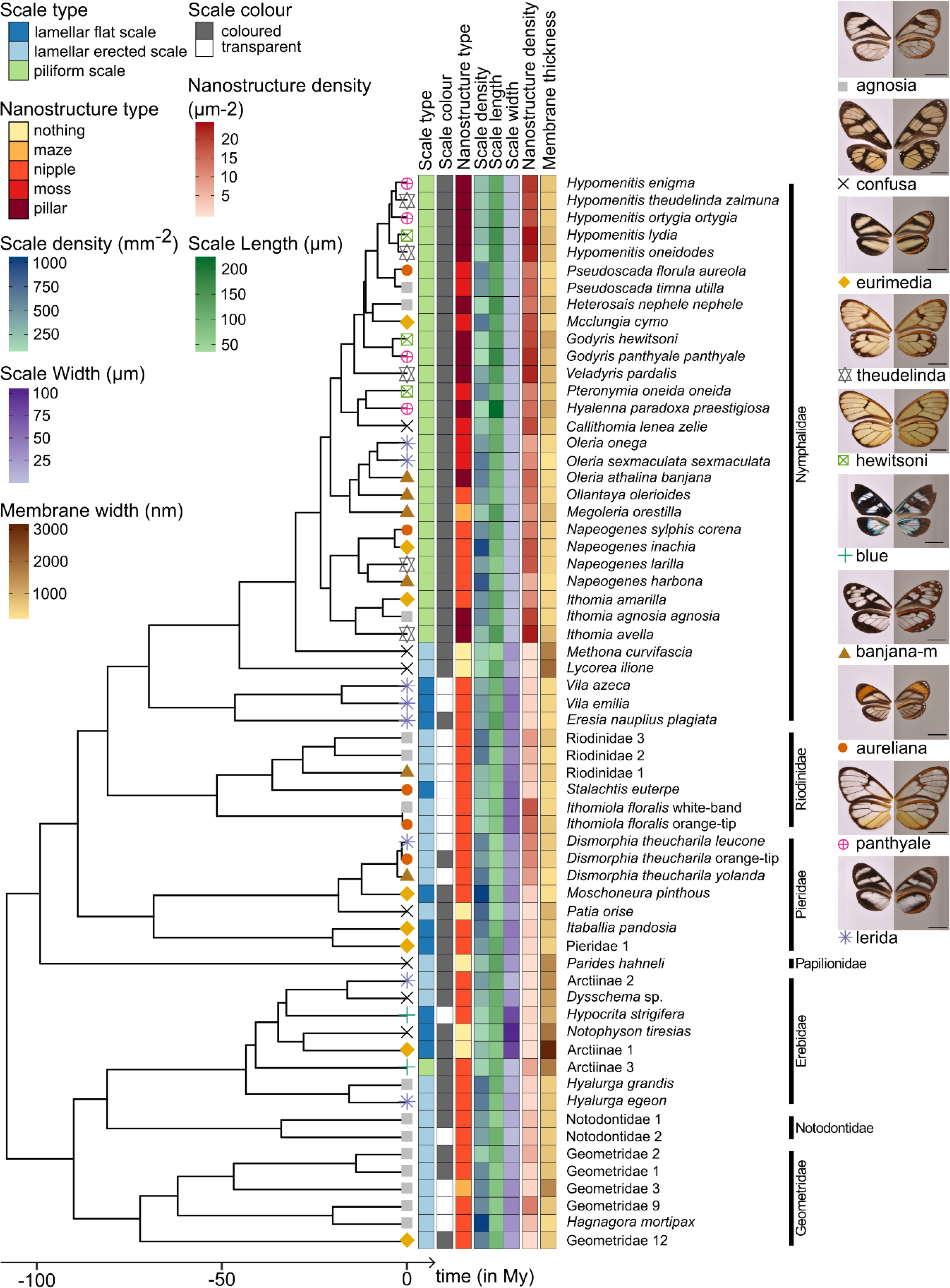
Phylogeny of the 62 species considered in this study and distribution of traits along the phylogeny. Mimicry rings are represented by a symbol and a specimen is given as an example for each mimicry ring. Dorsal side of wings has been photographed on a white background (left column) and ventral side on a gray background to highlight the transparent patches (right column). The x axis represents time in million years (My).

We also revealed an unexpected diversity of the nanostructures that cover wing membrane (Figure 2B). In our sample, we found five types of nanostructures: absent (10% of species), maze (3%), nipple arrays (55%), pillars (21%), and sponge-like (11%). Phylogenetic signal tests show that both micro- and nanostructure features are highly conserved in the phylogeny (Figure 3, Supplementary table 4), suggesting the existence of constraints in the developmental pathways underlying micro- and nanostructures. However, the value of δ, the metrics used to quantify phylogenetic signal of traits with discrete states (Borges et al., 2019), is higher for scale type than for nanostructures. This means that the phylogenetic signal is stronger for scale type than for nanostructures. Moreover, in the nymphalid tribe Ithomiini, which is highly represented in our dataset, microstructures seem to be more conserved (all species but the basal species *M. curvifascia* have piliform scales in transparent patches) than nanostructures (all five types of nanostructures, mixed in the Ithomiini clade, Figure 3).

We then investigated the convergence of structures among co-mimetic species by testing whether co-mimetic species shared structures more often than expected at random and given the phylogeny (see methods for details). We show that, across all mimicry rings, co-mimetic species share structural features (either scale type, nanostructures type or *structural syndrome*, defined as the association of scale type and nanostructure type) more than expected at random and given the phylogeny (Figure 4). The fact that the test remains significant when phylogenetic correction is applied means that structural features are convergent between co-mimetic species. We considered convergence of structural features for each individual mimicry ring (figure 4D and Supplementary table 3) and we showed that microstructures are convergent for ‘agnosia’ and ‘confusa’ mimicry rings, where species mainly have erected scales. In other mimicry rings (‘hewitsoni’, ‘panthyale’, ‘theudelinda’), species all have similar piliform scales but this similarity is likely due to shared ancestry and not to convergence. Regarding nanostructural features, we revealed convergent evolution for ‘agnosia’, ‘lerida’, ‘eurimedia’, ‘panthyale’ and ‘theudelinda’ mimicry rings, characterized by nipples for the three first ones and by pillar for the two last ones (Figure 4D, Supplementary table 3). We showed convergence in structural syndromes (association between micro- and nanostructures) for ‘agnosia’, where 71% of species harbour a combination of erected scales and nipples and for ‘panthyale’ where 100% of species combine piliform scales and pillar (Figure 4D, Supplementary table 3). For the mimicry ring ‘theudelinda’, 80% of species harbour a combination of piliform scales and pillars, but this similarity is best explained by shared ancestry.

**Figure 4.**
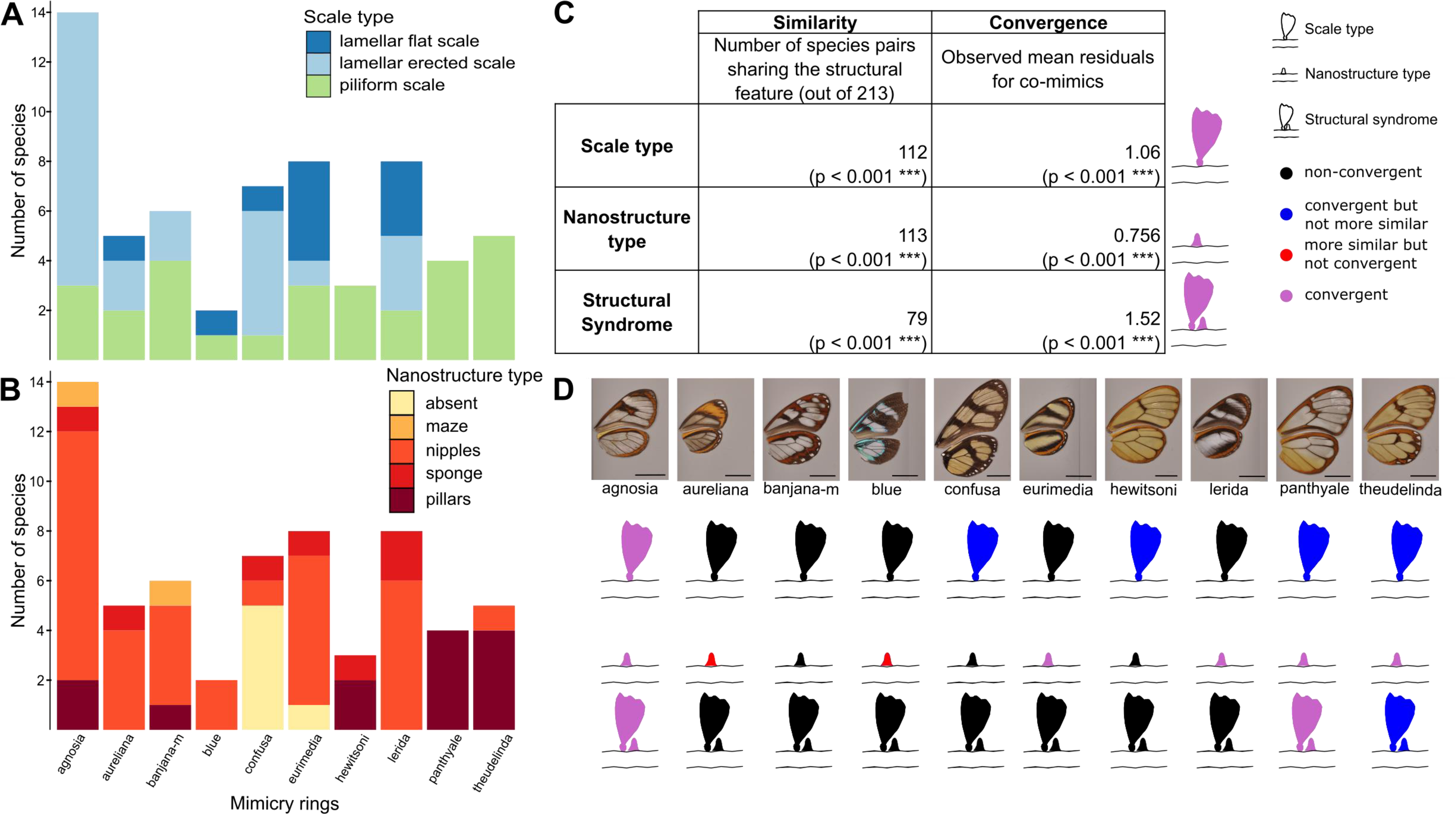
Convergence of structures underlying transparency. **A,B. Distribution of micro- (A) and nanostructures (B) among the different mimicry rings (indicated at the bottom on panel B).** **C. Results of the test of convergence for structural features (either scale type, nanostructure type or structural syndrome, i. e., the association of scale type and nanostructure type).** To test for similarity independently of the underlying process, we compare whether the number of co-mimetic species sharing the same structural feature (out of 213 co-mimetic species pairs) is higher than expected at random. To do so, we randomize 10000 times the sharing variable over all pairs of species and we calculate the p-value (indicated in brackets) as the proportion of randomisations where the number of co-mimetic species sharing the structural feature is higher than the observed number of co-mimetic species pairs sharing the structural feature. To test for convergence on structural features, we tested whether the observed mean residuals of the generalised linear model linking structure sharing and phylogenetic distance was higher than expected given the phylogeny and we calculated the p-value (indicated in brackets) as the proportion of randomisations of model’s residuals where the mean residuals for co-mimetic species is higher than the observed mean residuals for co-mimetic species. **D. Graphical representation of the results of the test of convergence for each mimicry ring.** For each mimicry ring, we tested whether the structural features were more similar than expected at random and given the phylogeny (with the same tests described above, see Supplementary table 3 for details). We represented the results for scale type, nanostructure type and structural syndrome. Black structures indicate neither more similar structures than expected at random nor convergent structures; red structures indicate structure more similar than expected at random but not convergence; blue structures indicate structures not more similar than expected at random but convergent and purple structures indicate convergent structures.

The fact that both transmission properties and underlying structures show some degree of convergence raises the question of whether specific structures have been selected in co-mimetic species because they confer a peculiar visual aspect, typical of the mimicry ring. To address this question, we investigated the link between structural features and transmission properties in transparent patches.

### Link between structural features and transmission properties

To investigate whether transmission properties depend on structural features we used the above measurements of the specular transmittance of transparent patches of each species and we calculated the mean transmittance over 300-700nm, hereafter called mean transmittance, for each spectrum. The physical property ‘mean transmittance’ (a proxy for the degree of transparency), is correlated to what is predicted to be perceived by predators based on vision modelling, (Supplementary result 1 for details). Across the 62 species, the mean transmittance ranges from 0.0284% in *Eresia nauplius* to 71.7% in *Godyris panthyale* (mean: 29.2%, median: 31.6%, Supplementary table 5). We performed Phylogenetic Generalized Least Squares (PGLS) to assess the relationship between mean transmittance and micro- and nanostructural features (type, insertion, colour, length, width and density of scales; type and density of nanostructures; wing membrane thickness; including some interactions), while accounting for the phylogeny. We retained as best models all models within 2 AICc units of the minimal AICc value. Following this procedure, eight models were retained (see below).

Mean transmittance depends mainly on scale type, scale density and nanostructure density, and to a lesser extent on membrane thickness and scale colour (Figure 5A, Supplementary table 6). The effect of scale type is retained in all eight models and is significant in all of them. Wings covered with piliform scales transmit more light than those covered with lamellar scales (Figure 5B). Among wings covered with lamellar scales, those with erected scales transmit more light than those with flat scales. The effect of scale density is retained in the eight best models and is significant in five of those (Supplementary table 6): mean transmittance decreases as scale density increases. The effect of nanostructure density is retained in six models and is significant in four of those: mean transmittance increases when nanostructure density increases (Figure 5C).

**Figure 5.**
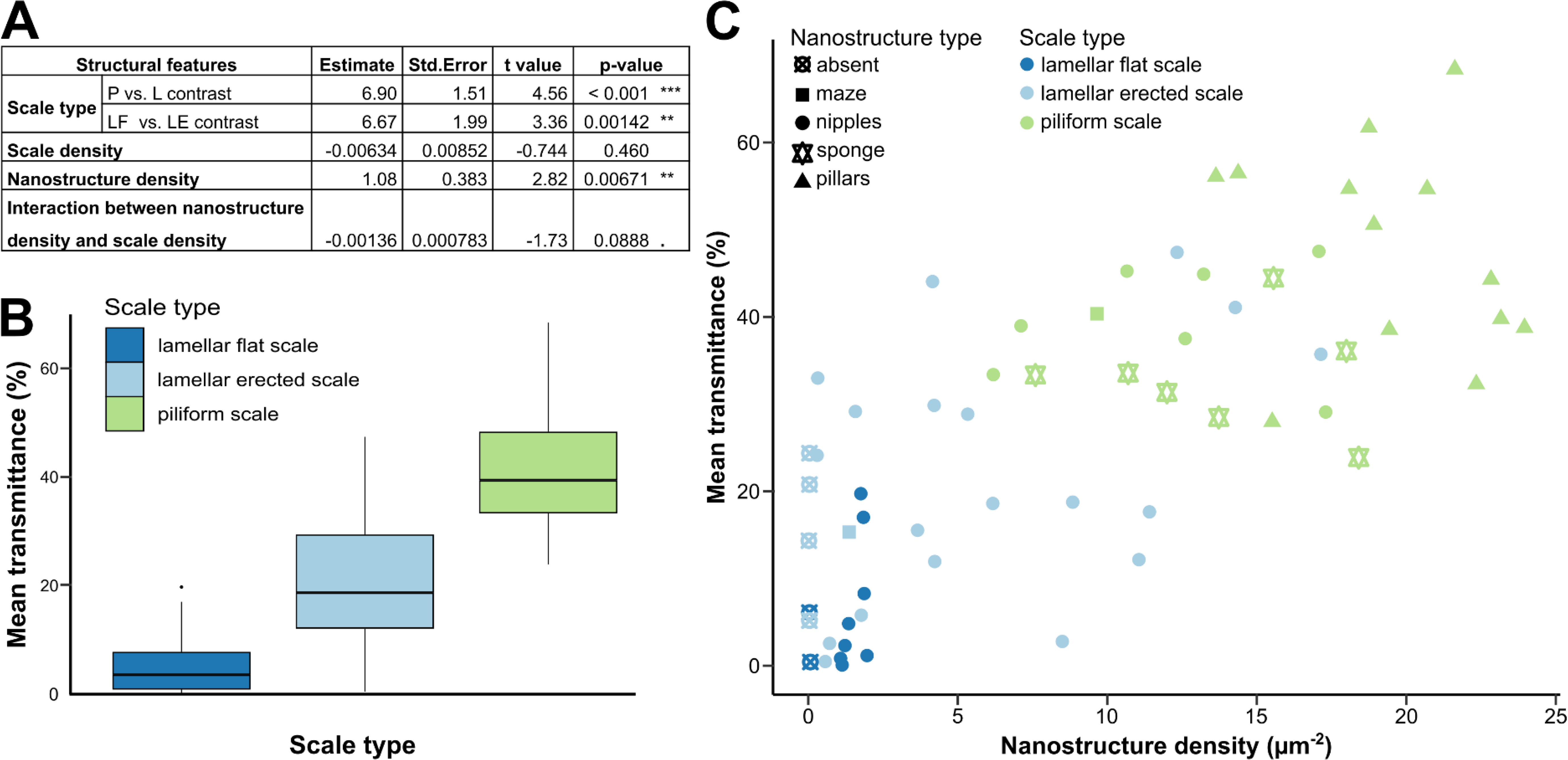
Link between mean transmittance over 300-700 nm and structural features. **A.** Results of the best PGLS model (F_5,56_ = 26.65 (p-value <0.001 ***), AICc = 469.9, R_adj_^2^ = 0.678, λ = 0 (p-value < 0.001 ***)) linking mean transmittance and micro- and nanostructure features. The explicative variables haven’t been scaled or centred. Nanostructure density has been measured in µm^-2^, Scale density in mm^-2^. Scale type is a categorical variable with 3 levels (either flat lamellar scale (LF), erected lamellar scale (LE) or piliform scale (P)) **B.** Link between mean transmittance measured between 300 and 700 nm and scale type. **C.** Link between mean transmittance measured between 300 and 700 nm and nanostructure density, nanostructure type (represented by different shape) and scale type (represented by different colours). NB. We considered the spot corresponding to the location of the SEM images for mean transmittance.

The interaction between scale density and nanostructure density is retained in three out of eight models and it is marginally significantly different from zero in two of those three models (Supplementary table 6). The coefficient is always negative, meaning that the increase in light transmission due to the increase in nanostructure density is not as strong when scale density is high than when scale density is low. This suggests that the contribution of nanostructures to transparency is stronger when scale density is low.

The effect of membrane thickness is retained in three out of eight models and is significantly different from zero in one of them: light transmission decreases when membrane thickness increases.

Transparent scales, which do not contain pigments, transmit more light than coloured ones, which contain pigments; a relationship which is retained in three out of eight models and is marginally significantly different from zero in one model (Supplementary table 6).

Other variables that were included in the model (scale length and width, nanostructure type, the interaction between scale type and scale density and the triple interaction between scale length, width and density) are not retained in any models (Supplementary table 6). These results suggest that those variables do not have any strong effect on transparency.

## Discussion

We conducted the first comparative study on transparent aposematic mimetic Lepidoptera to assess whether transparency is involved in the aposematic signal, to uncover the diversity in structures underlying transparency and to assess the link between transparency and structural features.

### Convergence of transmission properties

Based on bird vision modelling applied to light transmission measurements, we showed that predators see transparent patches of species belonging to the same mimicry ring as more similar than expected at random, and given the phylogeny. This shows that transparent patches in co-mimetic species are under selection for convergence, mirroring what has been shown for coloured patches in opaque species (Bybee et al., 2012; Llaurens et al., 2014; Su et al., 2015; Thurman & Seymoure, 2016). This convergent resemblance, which regards mainly the degree of transparency (a general term to refer to what extent a patch appears transparent), suggests that transparent patches might be part of the aposematic signal. Nevertheless, convergence in properties of transparent patches may also result from other selective processes. Transparency is also involved in crypsis (Arias, Elias, et al., 2020), even in aposematic prey (Arias et al., 2019; McClure et al., 2019), and the degree of transparency needed to achieve crypsis may depend on the ambient light (Arias, Barbut, et al., 2020; Johnsen & Widder, 1998). Specifically, in bright environments only highly transparent prey are cryptic, whereas in darker environments moderately transparent prey can be cryptic. In our case, as co-mimetic species share their habitat (Chazot et al., 2014) and microhabitat (Willmott et al., 2017), characterized by a specific ambient light, we cannot rule out that the similarity in the degree of transparency observed between co-mimetic species is the result of selection for crypsis rather than aposematism. Moreover, habitat-specific conditions, such as temperature or humidity, could also affect the evolution of transparent patches. In fact, the habitat and microhabitat shared by co-mimetic species is also characterized by a set of abiotic factors. Therefore, we cannot rule out that the observed convergence is driven by such abiotic factors (related to thermoregulation for example) instead of predation pressure. However, several studies on Ithomiini butterflies have shown that multiple mimicry rings may coexist in the same localities (Chazot et al., 2014; Elias et al., 2008; Willmott et al., 2017). For example, the species belonging to the mimicry rings ‘banjana-m’, ‘panthyale’, ‘hewitsoni’ and ‘theudelinda’ are all high-altitude species that are found in the same localities, and are therefore exposed to the same environmental conditions (ambient light, temperature, humidity). The fact that these species differ in transmission properties of their transparent patches (supplementary figure 1) suggests that the convergence observed is likely driven by Müllerian mimicry, and is not only the result of selection for crypsis or local adaptation to abiotic factors.

This study therefore challenges our vision of transparency, which might have evolved under multiple selective pressures in aposematic butterflies. Transparency has been shown to be involved in camouflage and to decrease detectability by predators (Arias, Elias, et al., 2020), even in aposematic species (Arias et al., 2019). Yet, our results suggest that transparent patches might also participate in the aposematic signal and that selection acts on the transmission properties of these patches, particularly on the degree of transparency, but also on chromatic aspects. Therefore, transparent aposematic Lepidoptera benefit from a double protection from predation, which can act at different distances (Barnett et al., 2018; Cuthill, 2019; Tullberg et al., 2005): transparent aposematic species are less detectable than opaque species (McClure et al., 2019), but when detected they may be recognized as unpalatable by experienced predators, due to their aposematic wing pattern, and spared by those predators.

### Structural features underlying transparency

#### Diversity of structures underlying transparency

We revealed an unexpected diversity of structures underlying transparency. Across the 62 species of the study, we found different microstructures in the transparent patches: transparent and coloured flat scales, transparent and coloured erected scales and piliform scale. Forked piliform scales have previously been reported in the highly transparent nymphalid species *Greta oto* (Binetti et al., 2009; Siddique et al., 2015), which belongs to the mimetic butterfly Ithomiini tribe. Erected scales (*i. e.,* with a non-zero angle between the scale basis and the wing membrane) have been previously reported in the riodinid *Chorinea sylphina* (Dushkina et al., 2017) and in the nymphalid *Parantica sita* (Perez Goodwyn et al., 2009). Here we describe, as Gomez et al. (2020, in review), some species with coloured erected scales that are completely perpendicular to the wing membrane, such as in the ithomiine *Methona curvifascia*. Transparent scales have already been reported in the opaque papilionid *Graphium sarpedon* (Stavenga et al., 2010) and as Gomez et al (2020, in review) we are describing them for the first time in transparent Lepidoptera. Other means of achieving transparency reported in the literature are not observed among our species (e. g., wing membrane devoid of scales, Yoshida et al. 1996). However, our study is restricted to mimetic transparent butterflies and, as such, spans a relatively small number of families. For a larger scale study that investigates thoroughly the different structures that might be involved in transparency in all known families comprising species with partially or totally transparent wings, see Gomez et al. (2020, in review). Scales in Lepidoptera are not only involved in colour patterns but also play a role in hydrophobicity. The scale modifications underlying transparency described in our study may impair the waterproofing properties of wings as shown by Perez Goodwyn et al. (2009): the wing of the translucent papilionid *Parnassius glacialis* are less hydrophobic than most Lepidoptera wings. Transparency may therefore come at a cost, especially for tropical Lepidoptera living in humid environments.

We also revealed an unexpected diversity of nanostructures covering the wing membrane, which we classify into five categories: absence of nanostructures, maze-like, nipple arrays, sponge-like and pillar-shaped nanostructures. While nipple arrays and pillars have previously been described on the wing of the sphingid *Cephonodes hylas* (Yoshida et al., 1997) and in the nymphalid *G. oto* (Binetti et al., 2009; Siddique et al., 2015), respectively, maze-like nanostructures have only been reported on the corneal surface of insect eyes (Blagodatski et al., 2015). Moreover, the sponge-like type of nanostructures is reported here for the first time. Those nanostructures can be related to the classification proposed by Blagodatski et al. (2015): pillars are a subcategory of nipple arrays, with higher and more densely packed nipples with an enlarged basis; sponge-like nanostructures are similar to dimples (holes embedded in a matrix), although with much bigger and more profound holes. Nipples, mazes and dimples have been found to be produced by Turing’s reaction-diffusion models, a solid framework that explains pattern formation in biology (Turing, 1952). While the principle of formation can be elegantly modelled, developmental studies are needed to understand the process by which nanostructures are laid on butterfly wing membrane (Pomerantz et al., 2020).

#### Link between structural features and transmission properties

The diversity of structures underlying transparency described above raises the question of whether these different structures confer different visual aspects. We indeed showed that mean transmittance over 300-700 nm, which is a proxy of the degree of transparency, depends on several structural features: scale type, scale density, nanostructure density, wing membrane thickness and scale colour. To summarize, mean transmittance increases when membrane coverage decreases, either due to reduced scale surface and/or scale density, because there is less material interacting (reflecting, diffusing, or absorbing) with light. Mean transmittance also increases when nanostructure density increases. Light transmission is indeed negatively correlated to light reflection and nanostructures are known to have anti-reflective properties, as demonstrated in the sphingid *Cephonodes hylas* (Yoshida et al., 1997) and in the nymphalid *G. oto* (Siddique et al., 2015). Reflection increases as the difference in refractive index between air and organic materials increases. Nanostructures create a gradient of refractive index between air and wing tissue, and gradient efficiency in reducing reflection increases with a smooth increase in proportion of chitin inside the nanostructures. For instance, pillars with conical bases are more effective at cancelling reflection than cylinders because cones produce a smoother air:chitin gradient from air to wing than cylinders (Siddique et al., 2015). Nanostructure shape is thus important in creating a smooth gradient. In our case, nanostructure density is highly correlated to nanostructure type, which we have defined according to their shape (phylogenetic ANOVA on nanostructure density with nanostructure type as factor: F = 26.26, p-value = 0.001, see supplementary result 2 for details). Specifically, the nanostructures whose shape likely creates the smoother gradient (pillar and moss) are also the denser ones. When nanostructure density increases, light reflection thus decreases. Light can either be transmitted, reflected or absorbed, and assuming that the chitin wing membrane only absorbs a small amount of light between 300 and 700 nm (Stavenga et al., 2014), when light reflection decreases because of the presence of nanostructures light transmission necessarily increases, which explains the positive effect of nanostructure density on mean transmittance.

We showed that mean transmittance decreases when membrane thickness increases, because wing membrane is mainly made of chitin and even if chitin absorbs a little amount of light (Stavenga et al., 2014), thicker membranes, which contain more chitin, absorb more light than thinner ones, thereby reducing light transmission.

We finally showed that wings covered with transparent scales transmit more light than wings covered with coloured scales. This is due to the presence of pigments, such as melanins or ommochromes commonly found in butterfly scales, which absorb some part of the light spectrum, thereby reducing light transmission.

Given the high structural diversity uncovered, future studies should thoroughly quantify the relative contributions of micro and nanostructures on the produced optical effects, notably on reflection in transparent patches which may encourage bio-inspired applications for transparent materials.

#### Selection on optical properties as a potential driver of the evolution of structures

We showed that transmission properties are convergent among co-mimetic species and that they depend on the underlying structural features, which confer peculiar visual aspects, raising the question of the putative convergence of structural features among co-mimetic species. We indeed showed that despite the high phylogenetic signal of structures underlying transparency that points to the existence of developmental constraints, both micro- and nanostructural features are convergent among co-mimetic species for some mimicry rings. Convergence is also detected for structural syndromes (i.e. association between micro- and nanostructures). Our data suggest that nanostructures are more labile than microstructures. Nanostructures could therefore evolve more readily in response to selection on the degree of transparency. We showed that the presence and higher densities of nanostructures increase mean transmittance when scale density is already low, thereby allowing fine-tuning of transparency. The interplay between scales and nanostructures can thus modulate the degree of transparency and the selective pressures on the transmission properties of transparent patches may select specific associations of structural features.

To conclude, this study reveals convergence of transparency features in aposematic mimetic Lepidoptera, which may be the result of selection by predators, likely through aposematism, even though transparent patches may also be under other local selection pressures such as selection for crypsis or adaptation to climatic conditions. Transparency entails strong structural modifications of scales that might impair other functions such as thermoregulation (Berthier, 2005), hydrophobicity (Perez Goodwyn et al., 2009) and perhaps mate signalling. Transparency may therefore come at a cost in those large-winged insects, which may explain why it is not pervasive among Lepidoptera.

## Materials & Methods

For further details about materials and methods see the Supplementary Materials & Methods section in the Supplementary Information.

### Material

In this study, we focus on 62 different species represented by 1 or 2 specimens collected with hand nets in understory forests in Peru and Ecuador, by ourselves and private collectors (Supplementary table 5). They belong to 7 different families (Nymphalidae, Riodinidae, Pieridae, Papilionidae, Erebidae, Notodontidae, Geometridae) and represent 10 different mimicry rings, following the classification used in Ithomiini: ‘agnosia’, ‘aureliana’, ‘banjana-m’, ‘confusa’, ‘eurimedia’, ‘hewitsoni’, ‘lerida’, ‘panthyale’, ‘theudelinda’ (Chazot et al., 2014; Willmott et al., 2017; Willmott & Mallet, 2004). In addition, we call ‘blue’ a mimicry ring that does not include Ithomiini species.

### Phylogeny

We used both published and *de novo* (see ‘Phylogeny’ section in SI for detailed protocol) sequences from one mitochondrial gene and seven nuclear genes, representing a total length of 7433 bp to infer a molecular phylogeny (knowing that for many taxa there are missing data, see Supplementary table 7). To improve the phylogeny topology, we added 35 species representing 8 additional families to the dataset (see Supplementary table 7 and SI). We performed a Bayesian inference of the phylogeny using BEAST 1.8.3. We forced the monophyly of some groups and we added eleven secondary calibration points (see Supplementary table 8) following Kawahara et al. (2019).

### Spectrophotometry

Specular transmittance was measured over 300-700 nm, a range of wavelengths to which both birds and butterflies are sensitive (Briscoe & Chittka, 2001; Hart, 2001) using a custom-built spectrophotometer (see ‘Spectrophotometry’ section in SI for details). For each species, we measured five different spots in the transparent patches on the ventral side of the forewing (see figure 1 for location). We computed mean transmittance over 300-700 nm from smoothed spectra using Pavo2 (Maia et al., 2019), as a proxy for transparency: wing transparency increases as mean transmittance increases. On a subset of 16 species, we measured 2 to 3 specimens per species and given that measurements were repeatable (see ‘Spectrophotometry’ section in SI), we retained only 1 specimen per species for optical measurements.

### High-resolution imaging and structure characterisation

We observed structures with a digital photonic microscope (Keyence VHX-5000) to determine scale form (lamellar scale vs. piliform scale), scale colour (coloured vs. transparent) and scale insertion (flat vs. erected) on ventral side. We defined as scale type the interaction between scale form and scale insertion (erected lamellar scale, flat lamellar scale and piliform scale). Wings were imaged using SEM (Zeiss Auriga 40) to determine nanostructure type and to measure scale density, scale length and width, membrane thickness, and nanostructure density (see SI for more details). We also determined for each species the structural syndrome, defined as the association between micro- and nanostructural features. On a subset of 3 species, we measured 10 specimens per species, each specimen being measured twice for density and five times for scale dimensions. Given that scale structural features were shown to be repeatable (see ‘High-resolution imaging and structure characterization’ section in SI) within species we retained one specimen per species in structure characterisation.

### Vision models

We used bird vision modelling on the smoothed transmission spectra to test whether transparent patches of co-mimetic species are perceived as similar by birds. Birds differ in their sensitivity to UV wavelength: some are more sensitive to UV (UVS vision) than others (VS vision). As predators of neotropical butterflies can belong to either category (Dell’Aglio et al., 2018), we used wedge-tailed shearwater (*Puffinus pacificus*) as a model for VS vision (Hart, 2004) and blue tit (*Cyanistes caeruleus*) as model for UVS vision (Hart et al., 2000). We considered two different light environments differing in their intensity and spectral distribution: forest shade and large gap as defined by Endler (Endler, 1993; Doris Gomez & Théry, 2007). In our model, we considered that the butterfly was seen against the sky (light is just transmitted through the wing). We used the receptor-noise limited model of Vorobyev and Osorio (Vorobyev & Osorio, 1998) with neural noise and with the following relative cone densities 1:1.9:2.7:2.7 (for UVS:S:M:L, Hart et al. 2000) and 1:0.7:1:1.4 (for VS:S:M:L, Hart 2004) for UVS and VS vision respectively, and a Weber fraction of 0.1 for chromatic vision (Lind et al., 2014; Maier & Bowmaker, 1993) and 0.2 for achromatic vision (average of the two species studied in Lind, Karlsson, and Kelber 2013) for both visual systems to compute chromatic and achromatic contrasts. In total we calculated 4 different vision models, using the R package Pavo 2 (Maia et al., 2019), representing all combinations of bird visual systems and light environments.

We extracted the chromatic and achromatic contrasts between each pair of species in the dataset, comparing only analogous spots (i.e. occupying the same position) on the forewing.

### Statistical analyses

All statistical analyses were performed with the software R version 3.6.2 and 4.0.3 (R Core Team, 2019).

#### Convergence on optical properties

To assess whether transparent patches, as perceived by predators, were more similar than expected at random, we calculated the mean phenotypic distance (either chromatic or achromatic contrast) for co-mimetic species and we compared this mean phenotypic distance to a null distribution of this mean distance, where the phenotypic distance has been randomised 10000 times over the 1891 possible pairs of species, irrespective of their phylogenetic relationship. The p-value was calculated as the proportion of randomisations when the calculated mean distance for co-mimetic species was smaller than the observed mean distance. The result of this test allows us to determine whether co-mimetic species are perceived as similar by their main predators, irrespectively of the evolutionary underlying mechanism, which can be either shared ancestry of convergent evolution. To disentangle the two possible mechanisms, we accounted for the phylogenetic relationship between species by performing a linear regression between phenotypic distances and phylogenetic distances for each pair of species, following Elias et al. (2008). Pairs of species below the regression line (with a negative residual) are phenotypically more similar than expected given the phylogeny. To test whether pairs of co-mimetic species were mostly below the regression line, we calculated the observed mean residuals for co-mimetic species and we compared it to a null distribution of mean residuals for co-mimetic species, where residuals have been randomised 10000 times over the 1891 possible pairs of species. The p-value was calculated as the proportion of randomisations where the calculated mean residuals was smaller than the observed mean residuals for co-mimetic species. We also tested for each mimicry ring whether co-mimetic species were perceived as more similar as expected at random and given the phylogeny by applying the tests described above as follows: we calculated mean phenotypic distance and mean residuals, respectively, for pairs of species belonging to the considered mimicry ring and compared these means to the random distribution of phenotypic distance and residuals, respectively, of the model restricted to the same number of observations (*i.e.* pair species) as in the mimicry ring considered.

#### Phylogenetic signal

To assess whether transmission properties and structural features were conserved in the phylogeny, we estimated the phylogenetic signal of each variable. For quantitative variable (mean transmittance, scale density, scale length, scale width, nanostructure density and membrane thickness), we calculated both Pagel’s λ (Pagel, 1999) and Blomberg’s K (Blomberg et al., 2003) implemented in the R package ‘phytools’ (Revell, 2012). For multicategorical variables (scale type and nanostructure type), we used the δ-statistic (Borges et al., 2019) and we compared it to the distribution of values of δ when the trait is randomised along the phylogeny to estimate whether the trait is randomly distributed along the phylogeny. Finally, for binary variables (scale colour), we used Fritz and Purvis’ D (Fritz & Purvis, 2010) implemented in the R package ‘caper’ (Orme et al., 2018).

#### Convergence on structures

We tested whether structural features (microstructures, i.e. scales and nanostructures, but also structural syndrome, *i.e.* the association between microstructures and nanostructures) are more similar between co-mimetic species as expected at random. To do so, we considered every pair of species in our dataset and we calculated the number of co-mimetic species sharing the same structural features. We compared this number to the null distribution of the number of species sharing the same structural features where the structural feature has been randomised 10000 times, a method similar to that used in Willmott and Mallet (2004). We calculated the p-value as the proportion of randomisations where the number of species sharing structures is higher than the observed number. To determine whether this sharing of structures was due to convergent evolution, we performed a generalized linear model with a binomial error distribution linking the variable for structure sharing (1 if species shared the same structure, 0 otherwise) with phylogenetic distance. We then calculated the mean residuals of the model for co-mimetic species and we compared it to a null distribution of mean residuals for co-mimetic species, where residuals have been randomised 10000 times. The p-value was given by the proportion of randomisation where the calculated mean residuals for co-mimetic species was higher than the observed mean residuals for co-mimetic species.

#### Link between transparency (mean transmittance) and structures

To assess the link between structural features and the degree of transparency we only used the spectrophotometric data of the points that correspond to the location of the SEM images (between 1 and 3 points per species) and we calculated the average of mean transmittance over 300-700 nm for each specimen. We tested the link between this average mean transmittance and all the structural features we measured (scale type, scale colour, scale density, scale length, scale width, nanostructure type, nanostructure density, membrane thickness and the following interactions: interaction between scale type and scale density, interaction between scale density and nanostructure density and the triple interaction between scale density, scale length and scale width), while controlling for phylogenetic relationships by performing Phylogenetic Generalised Least Square regression (PGLS) implemented in the R package ‘caper’ (Orme et al., 2018). We compared all possible models with all the structural variables, but we prevented some variables from being in the same model because they were highly correlated, using the R package ‘MuMIn’ (Barton, 2019). Among the 308 models, we selected the best models (difference in AICc inferior to 2). Eight such models were retained.

## Acknowledgments/funding

We thank Jonathan Pairraire and Céline Houssin for the help with photonic imaging and Josquin Gerber and Edgar Attivissimo for helping with spectroscopic measurements. We thank Benoit Vincent for the identification of some Arctiini specimens. We are thankful to the Institut de Physique du Globe de Paris (IPGP) for giving us access to SEM and to the Peruvian and Ecuadorian governmental authorities for collection permits (021C⁄C-2005-INRENA-IANP, 002-2015-SERFOR-DGGSPFFS, 373-2017-SERFOR-DGGSPFFS, 005-IC-FAU-DNBAPVS/MA, 019-IC-FAU-DNBAPVS/MA). This work was funded by Clearwing ANR project (ANR-16-CE02-0012), HFSP project on transparency (RGP0014/2016) and a France-Berkeley fund grant (FBF #2015-58).

---

**Supplementary figure 1.**
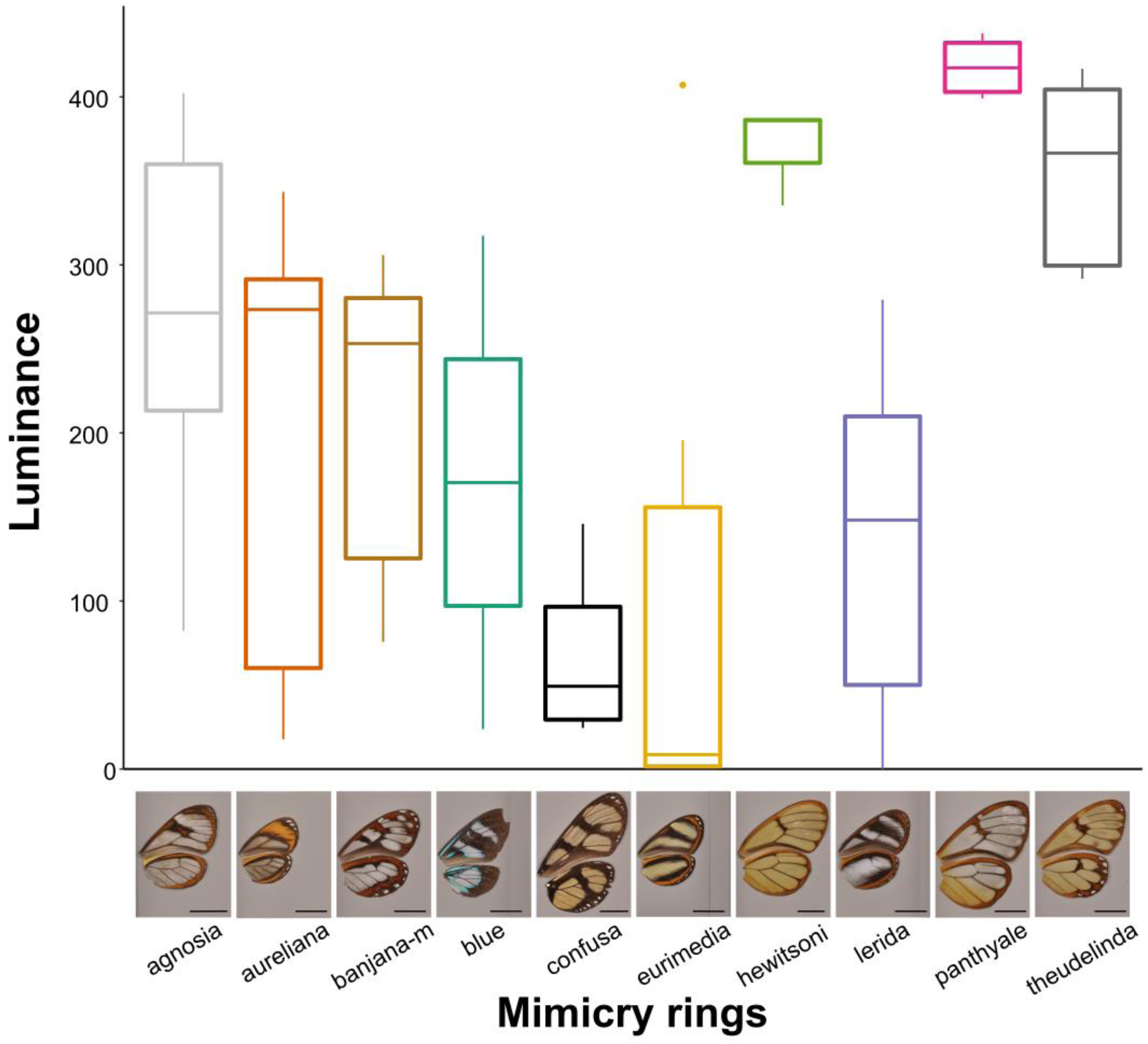
Differences in an avian analogue of luminance (quantum catch for brightness channel) between mimicry rings. The values presented here correspond to a vision model for large gap illuminants and UVS visual system, for the most proximal spot on the forewing. Results for other spots, other visual systems and illuminants are similar.

**Supplementary figure 2.**
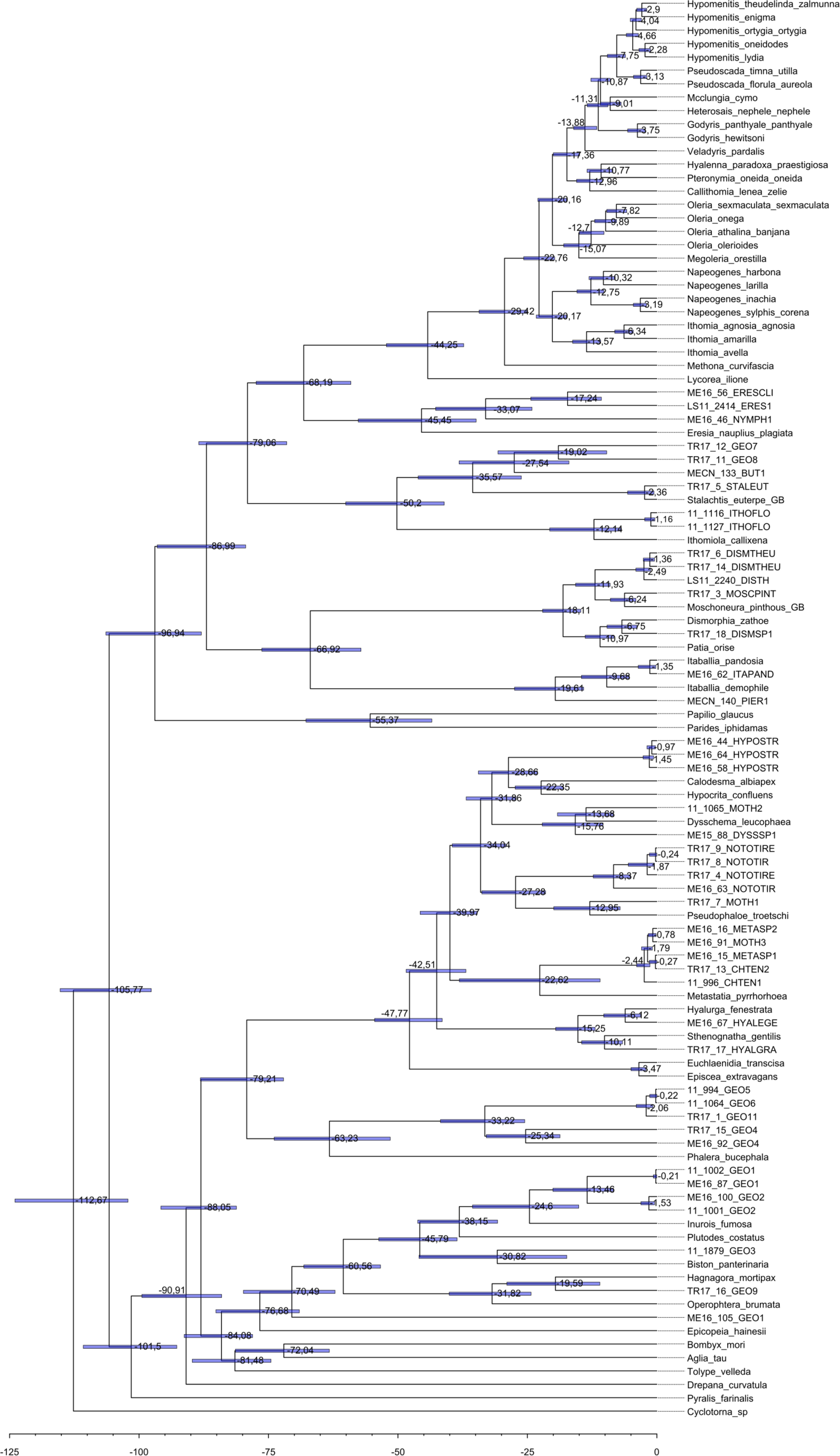
Maximum clade credibility tree of all the specimens obtained with BEAST 1.8.3. Each tip label represents one individual and the species name, when known, is given in the Supplementary table 7. Median node age is given at each node and the 95% interval of the node age post distribution is represented by a horizontal blue bar.

**Supplementary table 1a.** Tests of convergence of transparent patches, as perceived by predators, among co-mimetic species: achromatic contrasts. All the visual systems (VS and UVS) and the illuminants (large gap ‘lg’ and forest shade ‘fs’) tested are presented. We compare whether mean achromatic contrast (dL) between co-mimetic species is smaller than expected at random (expected mean dL for co-mimics ± standard deviation (sd)). To do so, we randomized the value of the achromatic contrast 10000 times over each pair of species and we calculated the p-value as the proportion of randomizations where the mean achromatic contrast for co-mimics is smaller than the observed mean achromatic contrast. We also considered whether co-mimetic species were more similar than what is expected according to their phylogenetic relationship. To do so, we did a linear model between achromatic contrasts and phylogenetic distances to account for the effect of phylogeny on the achromatic contrast and we considered the mean of residuals for co-mimetic species. If co-mimetic species are more similar than expected according to their phylogenetic relationship, the mean of residuals should be negative. To test whether the mean of residuals is smaller than expected according to the phylogeny, we randomized residuals over all pair of species and we calculated the mean of residuals for co-mimetic species. We calculated ‘p-value with phylogenetic correction’ as the proportion of randomizations where the mean of residuals is smaller than the observed mean of residuals. If p-value is smaller than 0.05, it means that co-mimetic species are more similar than expected at random and if p-value with phylogenetic correction is smaller than 0.05, it means that the observed similarity is due to convergence.

| Illuminant | Measured spot compared between 2 co-mimics | Avian visual model type | Similarity |  |  | Convergence |  |
| --- | --- | --- | --- | --- | --- | --- | --- |
| | | | Observed mean dL for co-mimics | Expected mean dL for co-mimics ± sd | p-value | Observed mean residuals for co-mimics | p-value with phylogenetic correction |
| fs | s1 | UVS | 6.779 | 7.412 ± 0.452 | 0.0819 (.) | -0.6670 | 0.0537 (.) |
| fs | s1 | VS | 6.768 | 7.406 ± 0.454 | 0.0792 (.) | -0.6646 | 0.0527 (.) |
| fs | s2 | UVS | 7.403 | 8.233 ± 0.540 | 0.0572 (.) | -0.8620 | **0.0365 (\*)** |
| fs | s2 | VS | 7.390 | 8.211 ± 0.536 | 0.0601 (.) | -0.8593 | **0.0455 (\*)** |
| fs | s3 | UVS | 5.150 | 6.310 ± 0.399 | **0.0009 (\*\*\*)** | -1.178 | **0.0002 (\*\*\*)** |
| fs | s3 | VS | 5.146 | 6.294 ± 0.400 | **0.001 (\*\*)** | -1.176 | **0.0005 (\*\*\*)** |
| fs | s4 | UVS | 5.148 | 6.727 ± 0.479 | **0.0002 (\*\*\*)** | -1.613 | **0.0001 (\*\*\*)** |
| fs | s4 | VS | 5.140 | 6.728 ± 0.491 | **0.0003 (\*\*\*)** | -1.608 | **< 0.0001 (\*\*\*)** |
| fs | s5 | UVS | 6.170 | 8.404 ± 0.601 | **< 0.0001 (\*\*\*)** | -2.265 | **< 0.0001 (\*\*\*)** |
| fs | s5 | VS | 6.163 | 8.396 ± 0.603 | **< 0.0001 (\*\*\*)** | -2.259 | **0.0001 (\*\*\*)** |
| lg | s1 | UVS | 6.778 | 7.408 ± 0.456 | 0.083 (.) | -0.6655 | 0.057 (.) |
| lg | s1 | VS | 6.765 | 7.396 ± 0.448 | 0.0793 (.) | -0.6625 | 0.0553 (.) |
| lg | s2 | UVS | 7.401 | 8.223 ± 0.538 | 0.0584 (.) | -0.8591 | **0.0388 (\*)** |
| lg | s2 | VS | 7.384 | 8.207 ± 0.538 | 0.057 (.) | -0.8559 | **0.0417 (\*)** |
| lg | s3 | UVS | 5.155 | 6.300 ± 0.404 | **0.001 (\*\*)** | -1.1755 | **0.0004 (\*\*\*)** |
| lg | s3 | VS | 5.140 | 6.293 ± 0.402 | **0.0005 (\*\*\*)** | -1.174 | **0.0005 (\*\*\*)** |
| lg | s4 | UVS | 5.147 | 6.740 ± 0.478 | **0.0001 (\*\*\*)** | -1.610 | **< 0.0001 (\*\*\*)** |
| lg | s4 | VS | 5.137 | 6.706 ± 0.478 | **0.0005 (\*\*\*)** | -1.604 | **0.0001 (\*\*\*)** |
| lg | s5 | UVS | 6.161 | 8.397 ± 0.595 | **< 0.0001 (\*\*\*)** | -2.262 | **< 0.0001 (\*\*\*)** |
| lg | s5 | VS | 6.153 | 8.364 ± 0.592 | **0.0001 (\*\*\*)** | -2.255 | **0.0002 (\*\*\*)** |

**Supplementary table 1b.** Tests of convergence of transparent patches, as perceived by predators, among co-mimetic species: chromatic contrasts. All the visual systems (VS and UVS) and the illuminants (large gap ‘lg’ and forest shade ‘fs’) tested are presented. We compare whether mean chromatic contrast (dS) between co-mimetic species is smaller than expected at random (expected mean dS for co-mimics ± standard deviation (sd)). To do so, we randomized the value of the chromatic contrast 10000 times over each pair of species and we calculated the p-value as the proportion of randomizations where the mean chromatic contrast for co-mimics is smaller than the observed mean chromatic contrast. We also considered whether co-mimetic species were more similar than what is expected according to their phylogenetic relationship. To do so, we did a linear model between chromatic contrasts and phylogenetic distances to account for the effect of phylogeny on the chromatic contrast and we considered the mean of residuals for co-mimetic species. If co-mimetic species are more similar than expected according to their phylogenetic relationship, the mean of residuals should be negative. To test whether the mean of residuals is smaller than expected according to the phylogeny, we randomized residuals over all pair of species and we calculated the mean of residuals for co-mimetic species. We calculated ‘p-value with phylogenetic correction’ as the proportion of randomizations where the mean of residuals is smaller than the observed mean of residuals. If p-value is smaller than 0.05, it means that co-mimetic species are more similar than expected at random and if p-value with phylogenetic correction is smaller than 0.05, it means that the observed similarity is due to convergence.

| Illuminant | Measured spot compared between 2 co-mimics | Avian visual model type | Similarity |  |  | Convergence |  |
| --- | --- | --- | --- | --- | --- | --- | --- |
| | | | Observed mean dS for co-mimics | Expected mean dS for co-mimics ± sd | p-value | Observed mean residuals for co-mimics | p-value with phylogenetic correction |
| fs | s1 | UVS | 2.727 | 2.833 ± 0.160 | 0.2565 | -0.1168 | 0.2278 |
| fs | s1 | VS | 2.520 | 2.621 ± 0.152 | 0.2534 | -0.1084 | 0.2329 |
| fs | s2 | UVS | 3.008 | 3.278 ± 0.188 | 0.075 (.) | -0.2777 | **0.049 (\*)** |
| fs | s2 | VS | 2.837 | 3.083 ± 0.167 | 0.0883 (.) | -0.2555 | 0.0709 (.) |
| fs | s3 | UVS | 2.861 | 3.071 ± 0.173 | 0.1124 | -0.2181 | 0.0985 (.) |
| fs | s3 | VS | 2.652 | 2.838 ± 0.167 | 0.1308 | -0.1929 | 0.1095 |
| fs | s4 | UVS | 2.703 | 2.879 ± 0.169 | 0.1509 | -0.1870 | 0.1204 |
| fs | s4 | VS | 2.497 | 2.672 ± 0.167 | 0.1519 | -0.1821 | 0.1282 |
| fs | s5 | UVS | 2.643 | 3.033 ± 0.178 | **0.0113 (\*)** | -0.3996 | **0.0084 (\*\*)** |
| fs | s5 | VS | 2.439 | 2.805 ± 0.171 | **0.0148 (\*)** | -0.3736 | **0.0107 (\*)** |
| lg | s1 | UVS | 2.707 | 2.816 ± 0.160 | 0.2422 | -0.1202 | 0.2137 |
| lg | s1 | VS | 2.520 | 2.624 ± 0.155 | 0.2563 | -0.1112 | 0.2263 |
| lg | s2 | UVS | 2.998 | 3.270 ± 0.184 | 0.0677 (.) | -0.2848 | 0.0503 (.) |
| lg | s2 | VS | 2.843 | 3.093 ± 0.182 | 0.0812 (.) | -0.2637 | 0.0652 (.) |
| lg | s3 | UVS | 2.846 | 3.057 ± 0.174 | 0.1137 | -0.2202 | 0.0944 (.) |
| lg | s3 | VS | 2.654 | 2.841 ± 0.168 | 0.1402 | -0.1938 | 0.1129 |
| lg | s4 | UVS | 2.679 | 2.867 ± 0.172 | 0.1441 | -0.1964 | 0.115 |
| lg | s4 | VS | 2.494 | 2.675 ± 0.170 | 0.1473 | -0.1898 | 0.1222 |
| lg | s5 | UVS | 2.616 | 3.015 ± 0.177 | 0.0099 (**) | -0.4084 | **0.0072 (\*\*)** |
| lg | s5 | VS | 2.425 | 2.798 ± 0.174 | 0.0124 (*) | -0.3816 | **0.0091 (\*\*)** |

**Supplementary table 2a.**
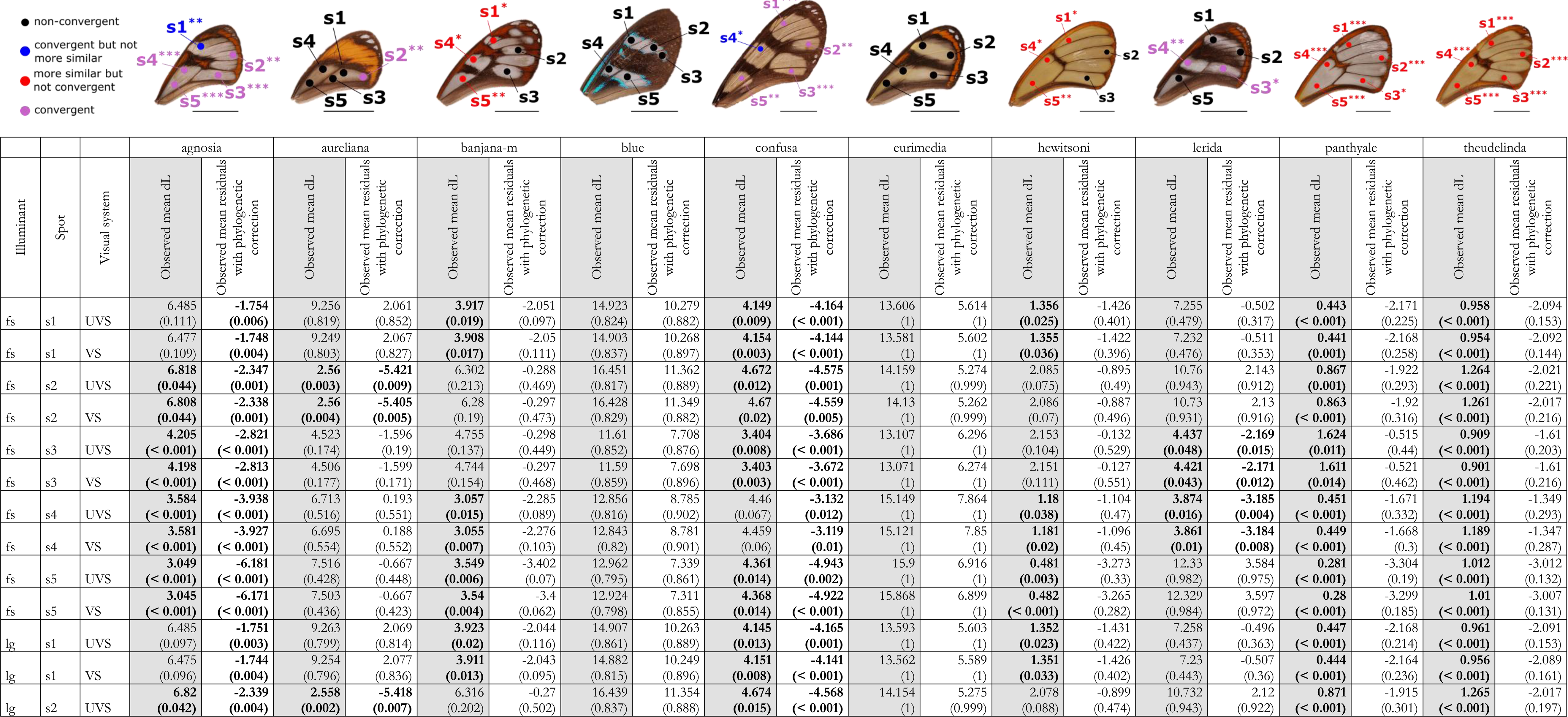
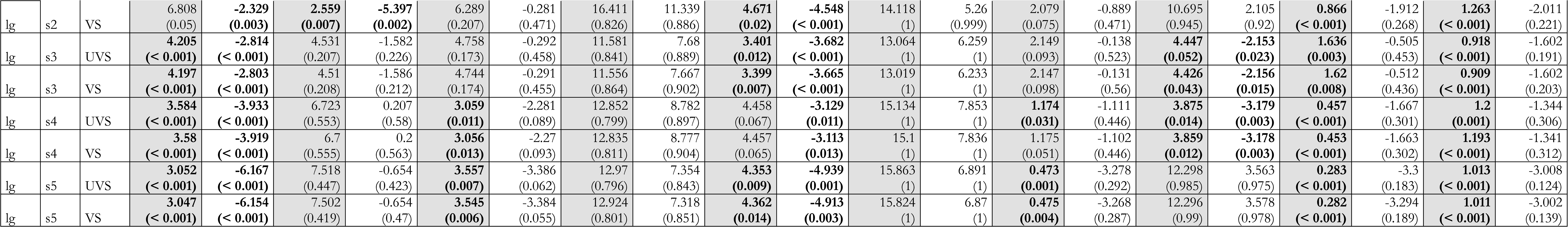
Results of the test of convergence of achromatic (dL) contrasts for each mimicry ring. All the visual systems (VS and UVS) and the illuminants (large gap ‘lg’ and forest shade ‘fs’) tested are presented. For each spot measured on the forewing and for each mimicry ring (see figure on top of the table), the mean achromatic contrast has been compared to what is expected at random (1^st^ column highlighted in grey for each mimicry ring, p-value in brackets) and to what is expected according to the phylogeny (2^nd^ column for each mimicry ring, p-value in brackets). Mean achromatic contrast statistically different from what is expected at random or given the phylogeny are highlighted in bold. To test whether mean achromatic contrast between co-mimetic species of a given mimicry ring is smaller to what is expected at random, we randomized the value of the achromatic contrast 10000 times over each pair of species, we calculated the mean achromatic contrast for these co-mimetic species for each randomisation and we calculated the p-value as the proportion of randomisations where the mean achromatic contrast for co-mimics is smaller than the observed mean achromatic contrast. To test whether the mean achromatic contrast between co-mimetic species of a given mimicry ring is smaller than what might be expected given their phylogenetic relationship, we did a linear model between achromatic contrasts and phylogenetic distances to account for the effect of phylogeny on the achromatic contrast and we considered the mean of residuals for co-mimetic species of a given mimicry ring. If co-mimetic species are more similar than expected according to the phylogeny, the mean of residuals should be negative. To test whether the mean of residuals is smaller than expected according to the phylogeny, we randomised residuals over all pair of species and we calculated the mean of residuals for co-mimetic species of a given mimicry ring. We calculated ‘p-value with phylogenetic correction’ as the proportion of randomisations where the mean of residuals is smaller than the observed mean of residuals. The figure above the table is a graphical representation of the results in the table (irrespective of the visual system or the illuminant as the results are similar): each spot measured is localised on the forewing and the results of the test are represented by the colour of the spot. Black spots stand for spots that are neither more similar than expected at random nor convergent (more similar when phylogenetic relationships are accounted for) between co-mimetic species. Blue spots stand for spots that are not more similar than expected at random but which are convergent because they are more similar that what might be expected according to their phylogenetic distance. Red spots stand for spots that are more similar than expected at random but not convergent. Purple spots stand for spots that are both more similar than expected at random and convergent (more similar than what might be expected based on phylogenetic distance). Tests’ p-values are represented with the following symbols: ‘***’ p < 0.001, ‘**’ p < 0.01, ‘*’ p < 0.05.

**Supplementary table 2b.** Results of the test of convergence of chromatic (dS) contrasts for each mimicry ring. All the visual systems (VS and UVS) and the illuminants (large gap ‘lg’ and forest shade ‘fs’) tested are presented. For each spot measured on the forewing and for each mimicry ring (see figure on top of the table), the mean chromatic contrast has been compared to what is expected at random (1^st^ column highlighted in grey for each mimicry ring, p-value in brackets) and to what is expected according to the phylogeny (2^nd^ column for each mimicry ring, p-value in brackets). Mean chromatic contrast statistically different from what is expected at random or given the phylogeny are highlighted in bold. To test whether mean chromatic contrast between co-mimetic species of a given mimicry ring is smaller to what is expected at random, we randomized the value of the chromatic contrast 10000 times over each pair of species, we calculated the mean chromatic contrast for these co-mimetic species for each randomisation and we calculated the p-value as the proportion of randomisations where the mean chromatic contrast for co-mimics is smaller than the observed mean chromatic contrast. To test whether the mean chromatic contrast between co-mimetic species of a given mimicry ring is smaller than what might be expected given their phylogenetic relationship, we did a linear model between chromatic contrasts and phylogenetic distances to account for the effect of phylogeny on the chromatic contrast and we considered the mean of residuals for co-mimetic species of a given mimicry ring. If co-mimetic species are more similar than expected according to the phylogeny, the mean of residuals should be negative. To test whether the mean of residuals is smaller than expected according to the phylogeny, we randomised residuals over all pair of species and we calculated the mean of residuals for co-mimetic species of a given mimicry ring. We calculated ‘p-value with phylogenetic correction’ as the proportion of randomisations where the mean of residuals is smaller than the observed mean of residuals. The figure above the table is a graphical representation of the results in the table (irrespective of the visual system or the illuminant as the results are similar): each spot measured is localised on the forewing and the results of the test are represented by the colour of the spot. Black spots stand for spots that are neither more similar than expected at random nor convergent (more similar when phylogenetic relationships are accounted for) between co-mimetic species. Blue spots stand for spots that are not more similar than expected at random but which are convergent because they are more similar that what might be expected according to their phylogenetic distance. Red spots stand for spots that are more similar than expected at random but not convergent. Purple spots stand for spots that are both more similar than expected at random and convergent (more similar than what might be expected based on phylogenetic distance). Tests’ p-values are represented with the following symbols: ‘***’ p < 0.001, ‘**’ p < 0.01, ‘*’ p < 0.05.

| Illuminant | Spot | Visual system | agnosia |  | aureliana |  | banjana-m |  | blue |  | confusa |  | eurimedia |  | hewitsoni |  | lerida |  | panthyale |  | theudelinda |  |
| --- | --- | --- | --- | --- | --- | --- | --- | --- | --- | --- | --- | --- | --- | --- | --- | --- | --- | --- | --- | --- | --- | --- |
|  |  |  | Observed mean dS | Observed mean residuals with phylogenetic correction | Observed mean dS | Observed mean residuals with phylogenetic correction | Observed mean dS | Observed mean residuals with phylogenetic correction | Observed mean dS | Observed mean residuals with phylogenetic correction | Observed mean dS | Observed mean residuals with phylogenetic correction | Observed mean dS | Observed mean residuals with phylogenetic correction | Observed mean dS | Observed mean residuals with phylogenetic correction | Observed mean dS | Observed mean residuals with phylogenetic correction | Observed mean dS | Observed mean residuals with phylogenetic correction | Observed mean dS | Observed mean residuals with phylogenetic correction |
| fs | s1 | UVS | 2.495<br>(0.068) | **-0.571**<br>**(0.008)** | **1.216**<br>**(0.012)** | **-1.557**<br>**(0.009)** | 1.825<br>(0.05) | -0.604<br>(0.173) | 2.653<br>(0.618) | 0.595<br>(0.651) | 2.51<br>(0.294) | -0.576<br>(0.116) | 4.401<br>(1) | 1.404<br>(0.998) | **0.362**<br>**(0.001)** | -1.175<br>(0.216) | 4.274<br>(0.998) | 1.343<br>(0.999) | **0.704**<br>**(0.002)** | -0.786<br>(0.209) | **1.066**<br>**(0.003)** | -0.546<br>(0.263) |
| fs | s1 | VS | **2.211**<br>**(0.036)** | **-0.617**<br>**(0.001)** | **1.125**<br>**(0.013)** | **-1.44**<br>**(0.007)** | 1.763<br>(0.068) | -0.495<br>(0.211) | 1.523<br>(0.48) | -0.402<br>(0.561) | 2.395<br>(0.35) | -0.45<br>(0.18) | 4.118<br>(1) | 1.353<br>(0.999) | **0.267**<br>**(0.002)** | -1.191<br>(0.222) | 4.149<br>(1) | 1.443<br>(0.999) | **0.59**<br>**(0.001)** | -0.826<br>(0.174) | **1.023**<br>**(0.005)** | -0.503<br>(0.255) |
| fs | s2 | UVS | **2.111**<br>**(< 0.001)** | **-1.481**<br>**(< 0.001)** | **0.775**<br>**(< 0.001)** | **-2.414**<br>**(0.001)** | 3.157<br>(0.455) | 0.441<br>(0.757) | 3.257<br>(0.598) | 1.052<br>(0.672) | 3.002<br>(0.347) | -0.618<br>(0.147) | 6.14<br>(1) | 2.644<br>(1) | 1.168<br>(0.105) | -0.319<br>(0.463) | 4.893<br>(0.996) | 1.488<br>(0.997) | **1.34**<br>**(0.036)** | -0.082<br>(0.51) | **0.678**<br>**(< 0.001)** | -0.914<br>(0.129) |
| fs | s2 | VS | **1.886**<br>**(< 0.001)** | **-1.5**<br>**(< 0.001)** | **0.573**<br>**(< 0.001)** | **-2.428**<br>**(0.002)** | 3.06<br>(0.491) | 0.513<br>(0.782) | 1.786<br>(0.528) | -0.273<br>(0.557) | 2.861<br>(0.339) | -0.552<br>(0.169) | 6.183<br>(1) | 2.888<br>(1) | 0.998<br>(0.096) | -0.374<br>(0.449) | 4.632<br>(0.999) | 1.424<br>(0.999) | **1.189**<br>**(0.038)** | -0.122<br>(0.464) | **0.633**<br>**(< 0.001)** | -0.839<br>(0.175) |
| fs | s3 | UVS | **2.19**<br>**(0.001)** | **-1.13**<br>**(< 0.001)** | 2.754<br>(0.393) | -0.25<br>(0.394) | **1.936**<br>**(0.044)** | -0.695<br>(0.14) | 4.74<br>(0.749) | 2.51<br>(0.822) | 3.3<br>(0.682) | -0.043<br>(0.498) | 5.586<br>(1) | 2.341<br>(1) | **0.359**<br>**(< 0.001)** | -1.306<br>(0.186) | 3.269<br>(0.68) | 0.095<br>(0.611) | 2.79<br>(0.432) | 1.176<br>(0.853) | **1.383**<br>**(0.016)** | -0.364<br>(0.355) |
| fs | s3 | VS | **1.895**<br>**(< 0.001)** | **-1.178**<br>**(< 0.001)** | 2.69<br>(0.472) | -0.083<br>(0.472) | 1.838<br>(0.056) | -0.582<br>(0.179) | 2.843<br>(0.608) | 0.805<br>(0.685) | 3.218<br>(0.76) | 0.123<br>(0.599) | 5.526<br>(1) | 2.523<br>(1) | **0.31**<br>**(< 0.001)** | -1.193<br>(0.242) | 3.026<br>(0.642) | 0.092<br>(0.595) | 2.43<br>(0.375) | 0.975<br>(0.814) | **1.261**<br>**(0.008)** | -0.32<br>(0.355) |
| fs | s4 | UVS | **2.117**<br>**(0.001)** | **-1.024**<br>**(< 0.001)** | 2.751<br>(0.454) | -0.059<br>(0.458) | **0.733**<br>**(< 0.001)** | **-1.687**<br>**(< 0.001)** | 2.295<br>(0.588) | 0.295<br>(0.655) | 3.653<br>(0.897) | 0.488<br>(0.815) | 5.644<br>(1) | 2.581<br>(1) | **0.376**<br>**(0.001)** | -1.033<br>(0.268) | 3.221<br>(0.75) | 0.233<br>(0.72) | **0.881**<br>**(0.008)** | -0.475<br>(0.335) | **1.083**<br>**(0.003)** | -0.411<br>(0.315) |
| fs | s4 | VS | **1.806**<br>**(< 0.001)** | **-1.105**<br>**(< 0.001)** | 2.683<br>(0.537) | 0.078<br>(0.602) | **0.6**<br>**(< 0.001)** | **-1.646**<br>**(0.001)** | 1.185<br>(0.374) | -0.673<br>(0.522) | 3.529<br>(0.924) | 0.596<br>(0.849) | 5.552<br>(1) | 2.713<br>(1) | **0.328**<br>**(0.002)** | -0.985<br>(0.29) | 3.047<br>(0.771) | 0.277<br>(0.703) | **0.694**<br>**(0.003)** | -0.57<br>(0.313) | **1.04**<br>**(0.012)** | -0.351<br>(0.34) |
| fs | s5 | UVS | **1.592**<br>**(< 0.001)** | **-1.694**<br>**(< 0.001)** | 2.444<br>(0.261) | -0.521<br>(0.255) | **1.39**<br>**(0.004)** | **-1.2**<br>**(0.024)** | 3.716<br>(0.659) | 1.532<br>(0.743) | 3.31<br>(0.702) | 0.002<br>(0.528) | 5.895<br>(1) | 2.685<br>(1) | **0.735**<br>**(0.021)** | -0.878<br>(0.288) | 4.055<br>(0.978) | 0.918<br>(0.979) | **0.714**<br>**(0.002)** | -0.847<br>(0.227) | **1.445**<br>**(0.025)** | -0.25<br>(0.418) |
| fs | s5 | VS | **1.329**<br>**(< 0.001)** | **-1.708**<br>**(< 0.001)** | 2.397<br>(0.327) | -0.344<br>(0.352) | **1.309**<br>**(0.005)** | **-1.083**<br>**(0.039)** | 4.01<br>(0.701) | 1.994<br>(0.812) | 3.208<br>(0.759) | 0.149<br>(0.629) | 5.739<br>(1) | 2.771<br>(1) | **0.664**<br>**(0.043)** | -0.824<br>(0.348) | 3.68<br>(0.946) | 0.78<br>(0.952) | **0.655**<br>**(0.002)** | -0.785<br>(0.261) | **1.384**<br>**(0.023)** | -0.18<br>(0.439) |
| lg | s1 | UVS | 2.455<br>(0.072) | **-0.594**<br>**(0.006)** | **1.2**<br>**(0.003)** | **-1.556**<br>**(0.007)** | **1.822**<br>**(0.043)** | -0.592<br>(0.177) | 2.553<br>(0.611) | 0.509<br>(0.68) | 2.484<br>(0.287) | -0.586<br>(0.13) | 4.443<br>(0.998) | 1.463<br>(0.998) | **0.354**<br>**(0.002)** | -1.17<br>(0.21) | 4.255<br>(0.999) | 1.341<br>(0.999) | **0.677**<br>**(0.002)** | -0.801<br>(0.202) | **1.051**<br>**(0.004)** | -0.549<br>(0.242) |
| lg | s1 | VS | **2.189**<br>**(0.026)** | **-0.642**<br>**(0.003)** | **1.115**<br>**(0.013)** | **-1.453**<br>**(0.009)** | 1.772<br>(0.072) | -0.488<br>(0.212) | 1.542<br>(0.499) | -0.385<br>(0.541) | 2.377<br>(0.302) | -0.473<br>(0.177) | 4.195<br>(1) | 1.426<br>(0.999) | **0.259**<br>**(0.001)** | -1.2<br>(0.18) | 4.166<br>(0.998) | 1.456<br>(0.999) | **0.568**<br>**(< 0.001)** | -0.849<br>(0.18) | **1.018**<br>**(0.005)** | -0.51<br>(0.224) |
| lg | s2 | UVS | **2.079**<br>**(< 0.001)** | **-1.51**<br>**(< 0.001)** | **0.745**<br>**(< 0.001)** | **-2.441**<br>**(< 0.001)** | 3.147<br>(0.46) | 0.436<br>(0.729) | 3.12<br>(0.574) | 0.921<br>(0.675) | 2.969<br>(0.309) | -0.649<br>(0.128) | 6.223<br>(1) | 2.729<br>(1) | 1.136<br>(0.091) | -0.343<br>(0.431) | 4.894<br>(0.996) | 1.491<br>(0.998) | **1.311**<br>**(0.039)** | -0.103<br>(0.519) | **0.665**<br>**(< 0.001)** | -0.918<br>(0.133) |
| lg | s2 | VS | **1.869**<br>(**< 0.001**) | **-1.533**<br>(**< 0.001**) | **0.549**<br>(**< 0.001**) | **-2.464**<br>(**< 0.001**) | 3.074<br>(0.509) | 0.516<br>(0.777) | 1.791<br>(0.47) | -0.274<br>(0.554) | 2.84<br>(0.348) | -0.588<br>(0.16) | 6.318<br>(1) | 3.008<br>(1) | 0.973<br>(0.096) | -0.401<br>(0.44) | 4.62<br>(0.997) | 1.398<br>(0.998) | **1.166**<br>(**0.033**) | -0.145<br>(0.498) | **0.626**<br>(**< 0.001**) | -0.847<br>(0.16) |
| lg | s3 | UVS | **2.14**<br>(**< 0.001**) | **-1.168**<br>(**< 0.001**) | 2.758<br>(0.348) | -0.232<br>(0.416) | **1.938**<br>(**0.044**) | -0.679<br>(0.154) | 4.537<br>(0.746) | 2.323<br>(0.831) | 3.283<br>(0.648) | -0.047<br>(0.503) | 5.722<br>(1) | 2.489<br>(1) | **0.362**<br>(**0.001**) | -1.284<br>(0.217) | 3.221<br>(0.644) | 0.06<br>(0.574) | 2.72<br>(0.411) | 1.125<br>(0.86) | **1.363**<br>(**0.009**) | -0.366<br>(0.369) |
| lg | s3 | VS | **1.859**<br>(**< 0.001**) | **-1.219**<br>(**< 0.001**) | 2.709<br>(0.466) | -0.066<br>(0.496) | 1.849<br>(0.053) | -0.571<br>(0.209) | 2.654<br>(0.595) | 0.619<br>(0.672) | 3.212<br>(0.776) | 0.113<br>(0.581) | 5.699<br>(1) | 2.693<br>(1) | **0.312**<br>(**0.002**) | -1.184<br>(0.174) | 2.999<br>(0.648) | 0.06<br>(0.572) | 2.376<br>(0.348) | 0.93<br>(0.818) | **1.247**<br>(**0.014**) | -0.326<br>(0.367) |
| lg | s4 | UVS | **2.058**<br>(**< 0.001**) | **-1.07**<br>(**< 0.001**) | 2.746<br>(0.471) | -0.05<br>(0.488) | **0.712**<br>(**< 0.001**) | **-1.694**<br>(**< 0.001**) | 2.169<br>(0.56) | 0.184<br>(0.67) | 3.624<br>(0.899) | 0.473<br>(0.795) | 5.752<br>(1) | 2.702<br>(1) | **0.364**<br>(**0.001**) | -1.03<br>(0.274) | 3.183<br>(0.766) | 0.208<br>(0.702) | **0.842**<br>(**0.009**) | -0.498<br>(0.335) | **1.07**<br>(**0.003**) | -0.409<br>(0.313) |
| lg | s4 | VS | **1.762**<br>(**< 0.001**) | **-1.156**<br>(**< 0.001**) | 2.696<br>(0.561) | 0.086<br>(0.55) | **0.589**<br>(**< 0.001**) | **-1.659**<br>(**0.001**) | 1.147<br>(0.394) | -0.711<br>(0.503) | 3.514<br>(0.911) | 0.575<br>(0.855) | 5.708<br>(1) | 2.863<br>(1) | **0.321**<br>(**0.003**) | -0.988<br>(0.317) | 3.037<br>(0.788) | 0.262<br>(0.743) | **0.664**<br>(**0.007**) | -0.596<br>(0.327) | **1.036**<br>(**0.008**) | -0.353<br>(0.34) |
| lg | s5 | UVS | **1.537**<br>(**< 0.001**) | **-1.732**<br>(**< 0.001**) | 2.461<br>(0.273) | -0.486<br>(0.276) | **1.379**<br>(**0.001**) | **-1.192**<br>(**0.025**) | 3.822<br>(0.665) | 1.66<br>(0.781) | 3.294<br>(0.699) | 0.003<br>(0.528) | 6.001<br>(1) | 2.808<br>(1) | **0.721**<br>(**0.039**) | -0.869<br>(0.313) | 3.95<br>(0.95) | 0.829<br>(0.957) | **0.695**<br>(**0.004**) | -0.843<br>(0.222) | **1.421**<br>(**0.012**) | -0.252<br>(0.409) |
| lg | s5 | VS | **1.288**<br>(**< 0.001**) | **-1.747**<br>(**< 0.001**) | 2.427<br>(0.349) | -0.308<br>(0.381) | **1.309**<br>(**0.002**) | **-1.075**<br>(**0.032**) | 4.16<br>(0.708) | 2.156<br>(0.833) | 3.207<br>(0.773) | 0.151<br>(0.602) | 5.884<br>(1) | 2.92<br>(1) | **0.653**<br>(**0.048**) | -0.817<br>(0.321) | 3.563<br>(0.92) | 0.667<br>(0.908) | **0.641**<br>(**0.001**) | -0.781<br>(0.233) | **1.37**<br>(**0.023**) | -0.178<br>(0.446) |

**Supplementary table 3.** Results of the test of convergence of structural features for each mimicry ring. We consider convergence in scale type (either piliform scale, lamellar erected scale or lamellar flat scale), in nanostructures type (either absent, maze, nipple, sponge or pillar) and in structural syndrome which corresponds to the association of one type of scale with one type of nanostructures. For each mimicry ring, we test the similarity of structures between co-mimetic species, independently of their phylogenetic relationship by comparing the sum of co-mimetic species sharing the same structure to what is expected at random (1^st^ column, highlighted in grey for each mimicry ring; we specified the number of co-mimetic species pair in brackets for each mimicry ring), based on 10000 randomisations of the variable ‘shared structures’. The p-value (in brackets) has been calculated as the proportion of randomisations where the sum of co-mimetic species sharing the same structure is higher than the observed sum of co-mimetic species sharing the same structure. We also tested the putative convergence of structures between co-mimetic species by performing a generalized linear model with binomial error linking the variable ‘shared structures’ (1 if the co-mimetic species share the same structures, 0 otherwise) and phylogenetic distance. We compared the mean of the model residuals for co-mimetic species to what is expected according to the phylogeny (2^nd^ column for each mimicry ring), based on 10000 randomisations of residuals along all pairs of species. The p-value (in brackets) was calculated as the proportion of randomisations where the mean of residuals in higher than the observed mean of residuals. The results are presented graphically in the figure above the table. Black structures indicate neither more similar than expected at random nor convergent structures; red structures indicate structure more similar than expected at random but not convergent; blue structures indicate structures not more similar than expected at random but convergent and purple structures indicate both more similar than expected at random and convergent structures.

|  | agnosia |  | aureliana |  | banjana-m |  | blue |  | confusa |  | eurimedia |  | hewitsoni |  | lerida |  | panthyale |  | theudelinda |  |
| --- | --- | --- | --- | --- | --- | --- | --- | --- | --- | --- | --- | --- | --- | --- | --- | --- | --- | --- | --- | --- |
|  | Sum of species sharing the same structure (out of 91) | Observed mean residuals with phylogenetic correction | Sum of species sharing the same structure (out of 10) | Observed mean residuals with phylogenetic correction | Sum of species sharing the same structure (out of 15) | Observed mean residuals with phylogenetic correction | Sum of species sharing the same structure (out of 1) | Observed mean residuals with phylogenetic correction | Sum of species sharing the same structure (out of 21) | Observed mean residuals with phylogenetic correction | Sum of species sharing the same structure (out of 28) | Observed mean residuals with phylogenetic correction | Sum of species sharing the same structure (out of 3) | Observed mean residuals with phylogenetic correction | Sum of species sharing the same structure (out of 28) | Observed mean residuals with phylogenetic correction | Sum of species sharing the same structure (out of 6) | Observed mean residuals with phylogenetic correction | Sum of species sharing the same structure (out of 10) | Observed mean residuals with phylogenetic correction |
| Scale type | 58<br>(**< 0.001**) | 2.114<br>(**< 0.001**) | 2<br>(0.783) | -0.760<br>(0.862) | 7<br>(0.151) | -0.028<br>(0.639) | 0<br>(0.388) | -2.941<br>(0.974) | 10<br>(0.121) | **1.306**<br>(**0.017**) | 9<br>(0.643) | 0.115<br>(0.527) | **3**<br>(**< 0.001**) | 1.221<br>(0.191) | 7<br>(0.862) | -0.357<br>(0.865) | **6**<br>(**< 0.001**) | 1.206<br>(0.145) | **10**<br>(**< 0.001**) | 1.253<br>(0.095) |
| Nanostructu<br>re type | 46<br>(**0.001**) | **0.631**<br>(**0.004**) | **6**<br>(**0.025**) | 1.06<br>(0.065) | 6<br>(0.267) | 0.201<br>(0.298) | **1**<br>(**< 0.001**) | 2.884<br>(0.075) | 10<br>(0.078) | 0.496<br>(0.144) | **15**<br>(**0.019**) | **0.767**<br>(**0.039**) | 1<br>(0.332) | -0.025<br>(0.3) | **16**<br>(**0.005**) | **0.927**<br>(**0.013**) | **6**<br>(**< 0.001**) | **2.951**<br>(**< 0.001**) | **6**<br>(**0.037**) | **1.157**<br>(**0.029**) |
| Structural<br>syndrom | 46<br>(**< 0.001**) | 3.032<br>(**< 0.001**) | 1<br>(0.548) | -0.397<br>(0.638) | 2<br>(0.495) | -0.416<br>(0.701) | 0<br>(0.179) | -1.324<br>(0.851) | 6<br>(0.054) | 1.153<br>(0.052) | 4<br>(0.556) | -0.371<br>(0.778) | 1<br>(0.076) | 0.056<br>(0.461) | 7<br>(0.082) | 0.435<br>(0.191) | 6<br>(**< 0.001**) | 3.117<br>(**0.024**) | 6<br>(**0.001**) | 1.373<br>(0.078) |

**Supplementary table 4.**
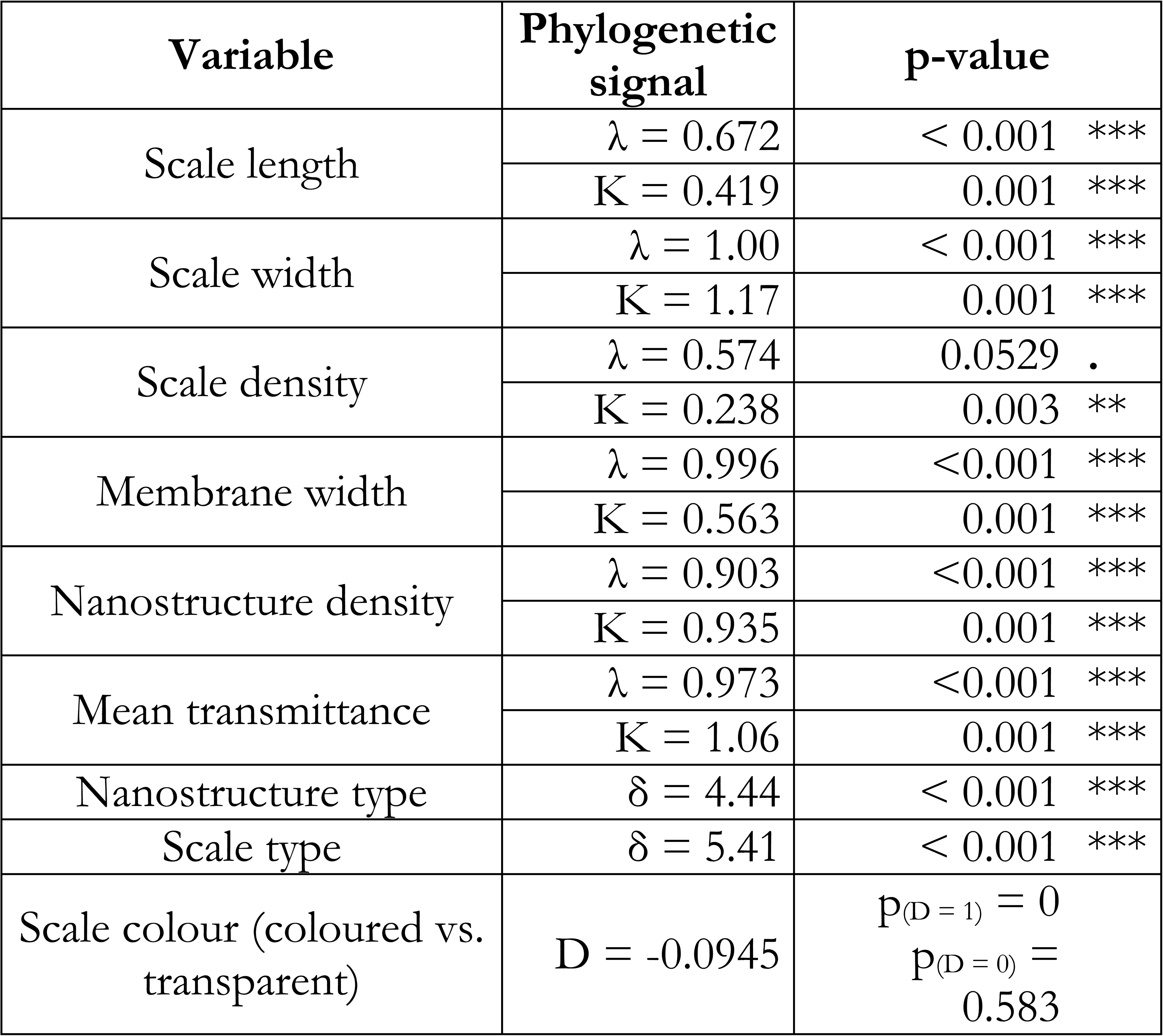
Phylogenetic signal for structural features and transmission properties. Measure of the phylogenetic signal (estimated as Pagel’s λ and Blomberg’s K for quantitative traits; δ for multicategorial traits and Purvis and Fritz’s D for binary traits) of the different features associated to micro- and nanostructures and of mean transmittance. When λ or K are equal to 0, the trait is distributed randomly across the phylogeny, whereas when λ or K are equal to 1 the trait evolves according to a Brownian motion model along the phylogeny. When D is equal to 1, the trait is randomly distributed across the phylogeny whereas when D is equal to 0, the trait evolves according to Brownian motion model along the phylogeny. The value of δ can be any positive real number and the higher this value, the higher the phylogenetic signal of the trait. For δ, to determine whether the distribution of the trait is different from a random distribution we randomized the trait 1000 times along the phylogeny, and we calculated δ for each randomisation. We then compared the value of δ to the distribution of values of δ under the random hypothesis and we calculated a p-value as the number of randomisations in which δ is higher than the value obtained for the real distribution of the trait.

**Supplementary table 5.**
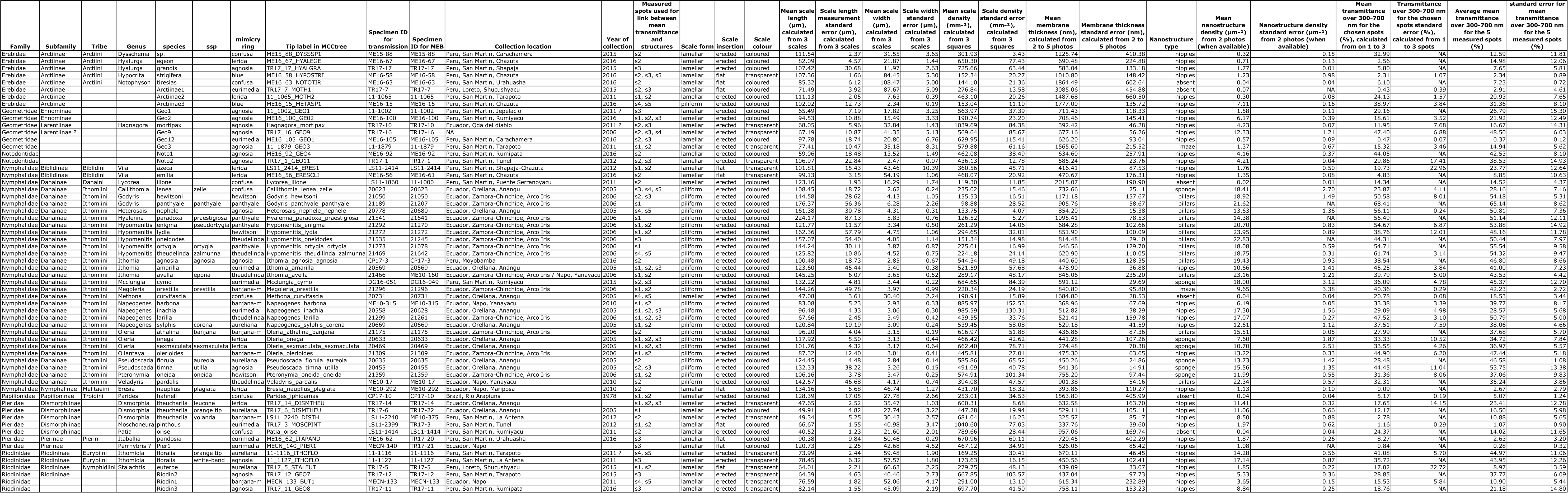
Information about specimens used for optical and structural measurements.

**Supplementary table 6.**
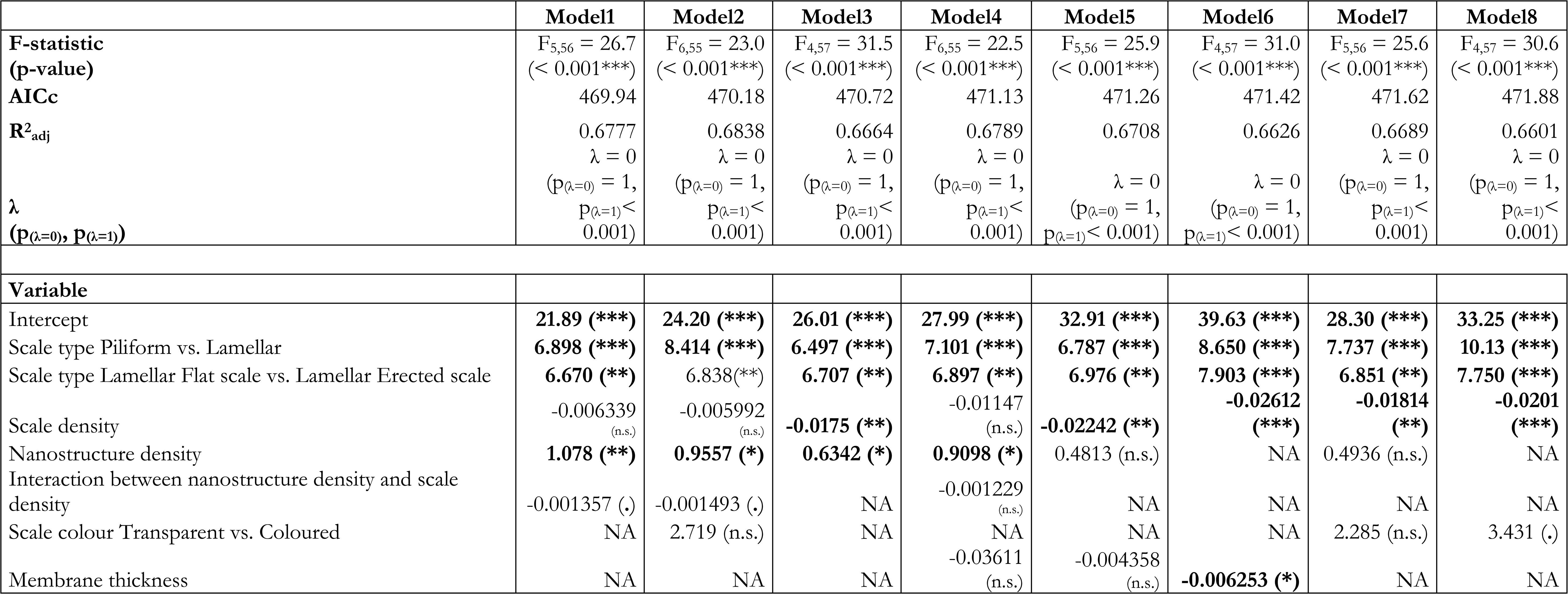
Results of the eight best PGLS (Phylogenetic Generalized Least Square) models (AICc within an interval of 2 of that of the best model). For each model, we give: the F statistic with the degrees of freedom in indices, the p-value of the model (in brackets), the corrected Akaike criterion (AICc) of the model, the adjusted R² and the value of lambda branch length transformation which has been estimated by maximum likelihood given the statistical model linking traits. When λ equals 1, the branch length of the phylogeny is unchanged, whereas when λ equals 0 branch length is set to zero, meaning that all species are considered independent. The ‘p-values’ for the value of λ, given in brackets, are the probability that λ is equal to 0 or to 1. We also give for each model the value of the coefficient estimate for each variable tested and the p-value (in brackets) is represented with the follow symbols: *** : p < 0.001; ** : p < 0.01; * : p<0.05;. : p < 0.1; n.s. : not significantly different from 0. NA means that the variable was not retained in the model

**Supplementary table 7.**
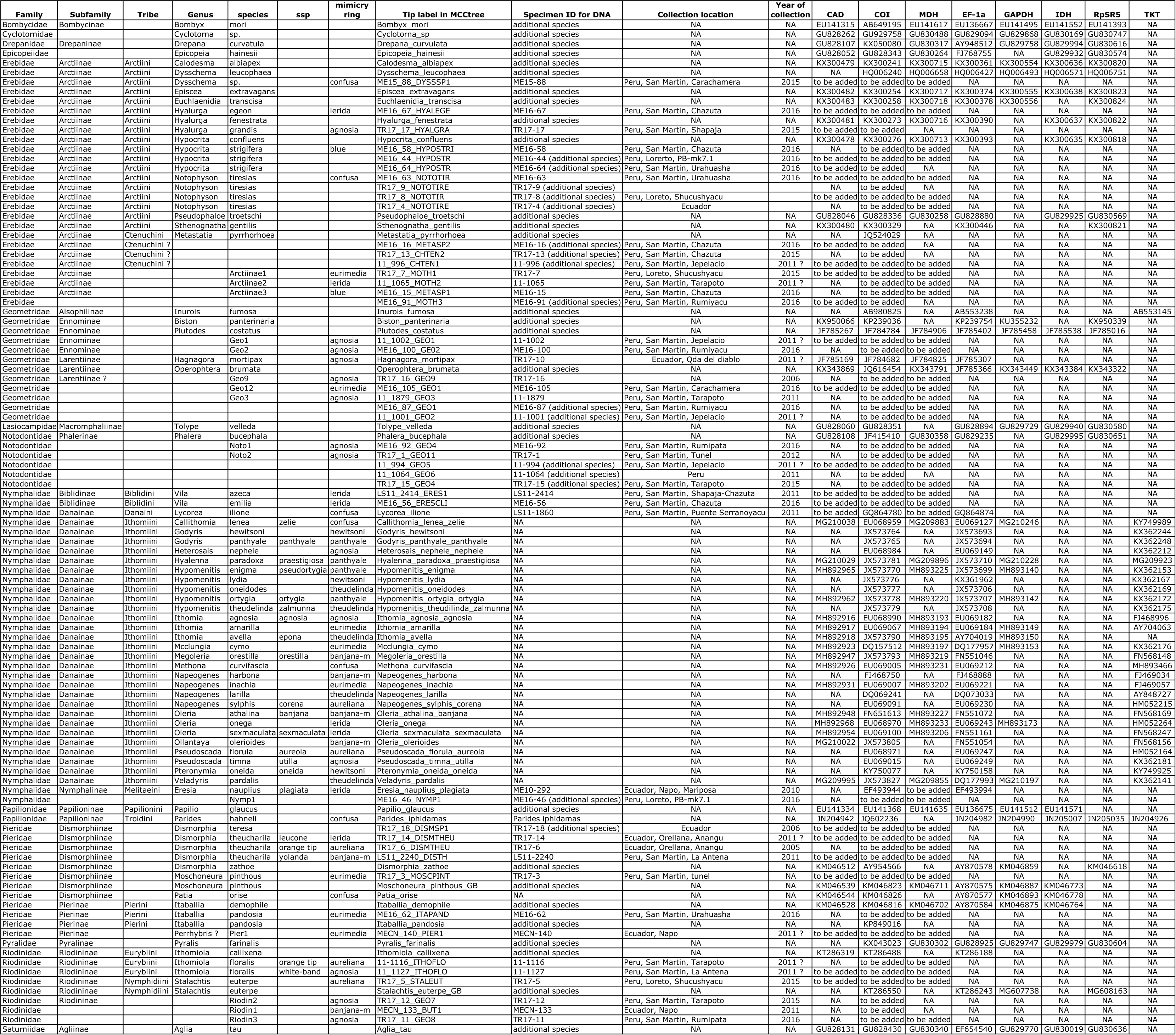
Information on specimens used to infer a phylogeny

**Supplementary table 8.**
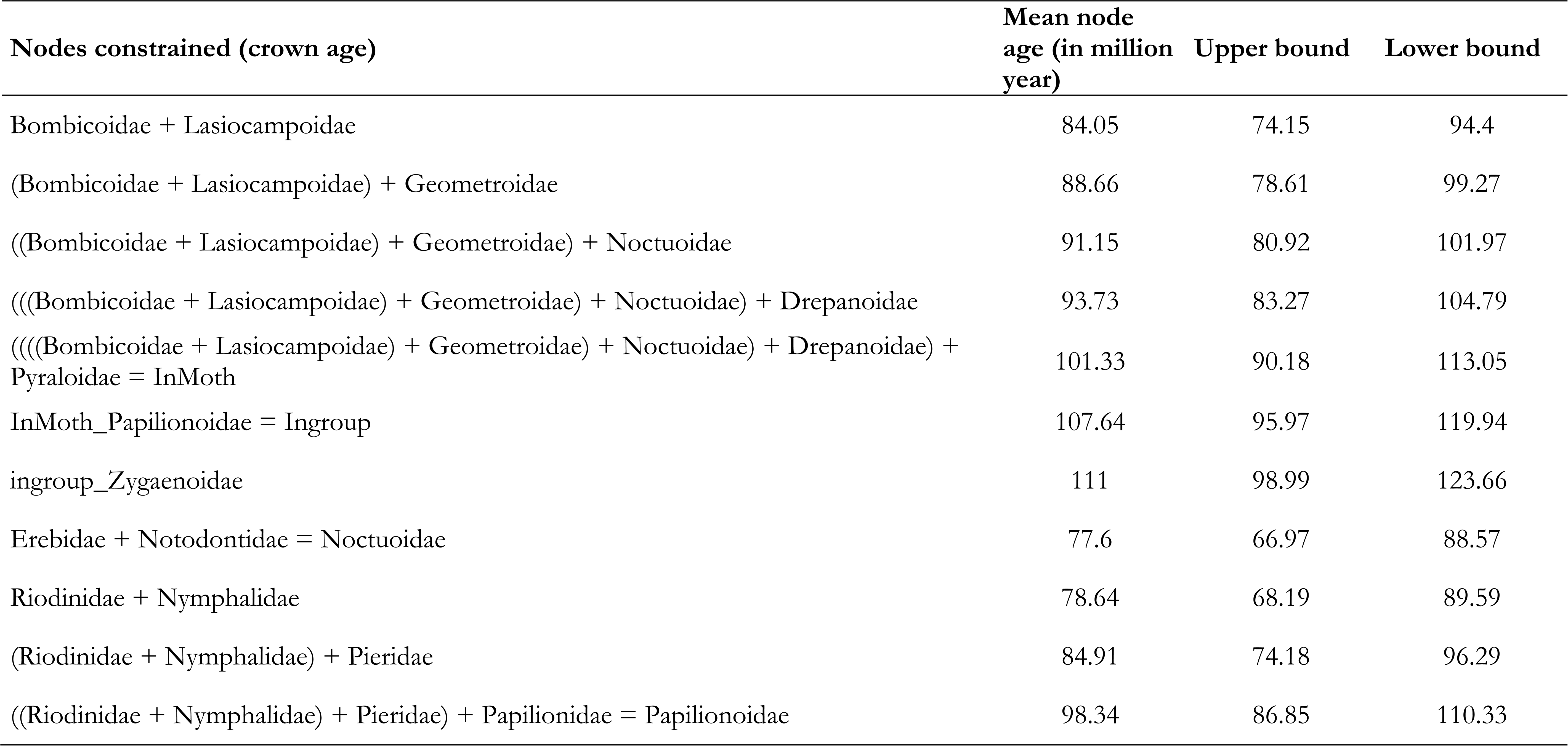
Node constraints used to calibrate the phylogeny. For all constraints, we used uniform distribution priors whose bound were determined according to 95% HSPD inferred by Kawahara et al. 2019 on their phylogeny of Lepidoptera.

**Supplementary table 9.** Results of the best partition (based on BIC) for the 8 different genes obtained with Partition Finder v1.0.1, with linked branch length and greedy algorithm. For each gene, pos1, pos2 and pos3 refer to codon positions. Only the substitution models available in BEAST were tested. GTR : general time reversible (base frequencies are variable, substitution matrix is symmetrical), HKY : Hasegawa-Kishino-Yano (base frequencies are variable, there are one transition rate and one tranversion rate), TrN : Tamura-Nei (base frequencies are variable, transversion rates are equal, transition rates are variable), I : proportion of invariable sites, G : gamma distribution (rate variation among sites is gamma distributed).

| Gene fragment and codon position | Best substitution model |
| --- | --- |
| CAD_pos1, IDH_pos1, MDH_pos1, TKT_pos1 | GTR+I+G |
| CAD_pos2, MDH_pos2, TKT_pos2 | GTR+I+G |
| CAD_pos3, MDH_pos3 | HKY+I+G |
| COI_pos1 | GTR+I+G |
| COI_pos2 | HKY+I+G |
| COI_pos3 | HKY+G |
| EF-1a_pos1, GAPDH_pos1, RpS5_pos1 | TrN+I+G |
| EF-1a_pos2, GAPDH_pos2, IDH_pos2, RpS5_pos2 | HKY+I+G |
| EF-1a_pos3 | GTR+I+G |
| GAPDH_pos3, IDH_pos3, RpS5_pos3 | GTR+I+G |
| TKT_pos3 | HKY+G |

**Supplementary table 10a.** Repeatability of transmission measurements and structural features. For each grouping factor (either the number of species or the number of individuals or the total number of different spots measured; indicated in the ‘number of groups’ column), we calculated the value of repeatability R based on several measurements of the same element of a grouping factor. The calculation of repeatability is based on mixed linear models. Confidence intervals are calculated with parametric bootstraping and p-values (associated to the test R > 0) are calculated with two methods : with likelihood ratio test comparing the likelihood of the model with and without the tested random effect and with permutation tests. We also calculated the coefficient of variation (CV, as the mean of the group devided by the standard error) for each group and we give here the median value of the CV distribution.

| Dependent Variable | Fixed effect | Random effect | Total number of observations | Number of groups | R | 95% confidence interval | p-value (obtained by LRT) | p-value (obtained by permutation) | Median CV (%) |
| --- | --- | --- | --- | --- | --- | --- | --- | --- | --- |
| Mean transmittance over 400-700 nm | | Species, Individual, spot | 476 obs | 11 groups<br>32 groups<br>160 groups | $R_{\text{species}} = \mathbf{0.586}$<br>$R_{\text{individual}} = 0.0535$<br>$R_{\text{spot}} = \mathbf{0.34}$ | [0.285 ; 0.752]<br>[~0 ; 0.155]<br>[0.201 ; 0.601] | < 0.001 ***<br>0.033 *<br>< 0.001 *** | 0.836<br>0.407<br>0.001 *** | 2.23 |
| Chroma over 400-700 nm | | Species, Individual, spot | 476 obs | 11 groups<br>32 groups<br>160 groups | $R_{\text{species}} = \mathbf{0.583}$<br>$R_{\text{individual}} = 0.0476$<br>$R_{\text{spot}} = \mathbf{0.355}$ | [0.271 ; 0.745]<br>[0 ; 0.161]<br>[0.210 ; 0.635] | < 0.001 ***<br>0.053 .<br>< 0.001 *** | 0.79<br>0.412<br>0.001 *** | 3.41 |
| Scale density | Species, sex | Individual, spot | 120 obs | 30 groups<br>60 groups | $R_{\text{individual}} = 0$<br>$R_{\text{spot}} = \mathbf{0.327}$ | [0 ; 0.18]<br>[0.144 ; 0.455] | 1<br>< 0.001 *** | 1<br>0.001 *** | 13.1 |
| Scale length |  |  |  |  |  |  |  |  |  |
| Bifid piliform scale | Species, sex | Individual, spot | 246 obs | 30 groups<br>50 group | $R_{\text{individual}} = 0.0779$<br>$R_{\text{spot}} = \mathbf{0.245}$ | [0 ; 0.259]<br>[0.0717 ; 0.406] | 0.196<br>< 0.001 *** | 0.318<br>0.001 *** | 7.69 |
| Monofid piliform scale | Species, sex | Individual, spot | 354 obs | 30 groups<br>60 groups | $R_{\text{individual}} = 0$<br>$R_{\text{spot}} = \mathbf{0.113}$ | [0 ; 0.0864]<br>[0.00917 ; 0.193] | 1<br><0.001 *** | 1<br>0.001 *** | 16.5 |
| Scale width |  |  |  |  |  |  |  |  |  |
| Bifid piliform scale | Species, sex | Individual, spot | 246 obs | 30 groups<br>50 groups | $R_{\text{individual}} = 0.149$<br>$R_{\text{spot}} = \mathbf{0.149}$ | [$2.31 \cdot 10^{-12}$ ; 0.315]<br>[0.0322 ; 0.292] | 0.026<br>< 0.001 *** | 0.166<br>0.001 *** | 8.65 |
| Monofid piliform scale | Species, sex | Individual, spot | 354 obs | 30 groups<br>60 groups | $R_{\text{individual}} = 0$<br>$R_{\text{spot}} = \mathbf{0.0816}$ | [0 ; 0.0599]<br>[0.0105 ; 0.131] | 1<br>< 0.001 *** | 1<br>0.001 *** | 9.33 |

**Supplementary table 10b.**
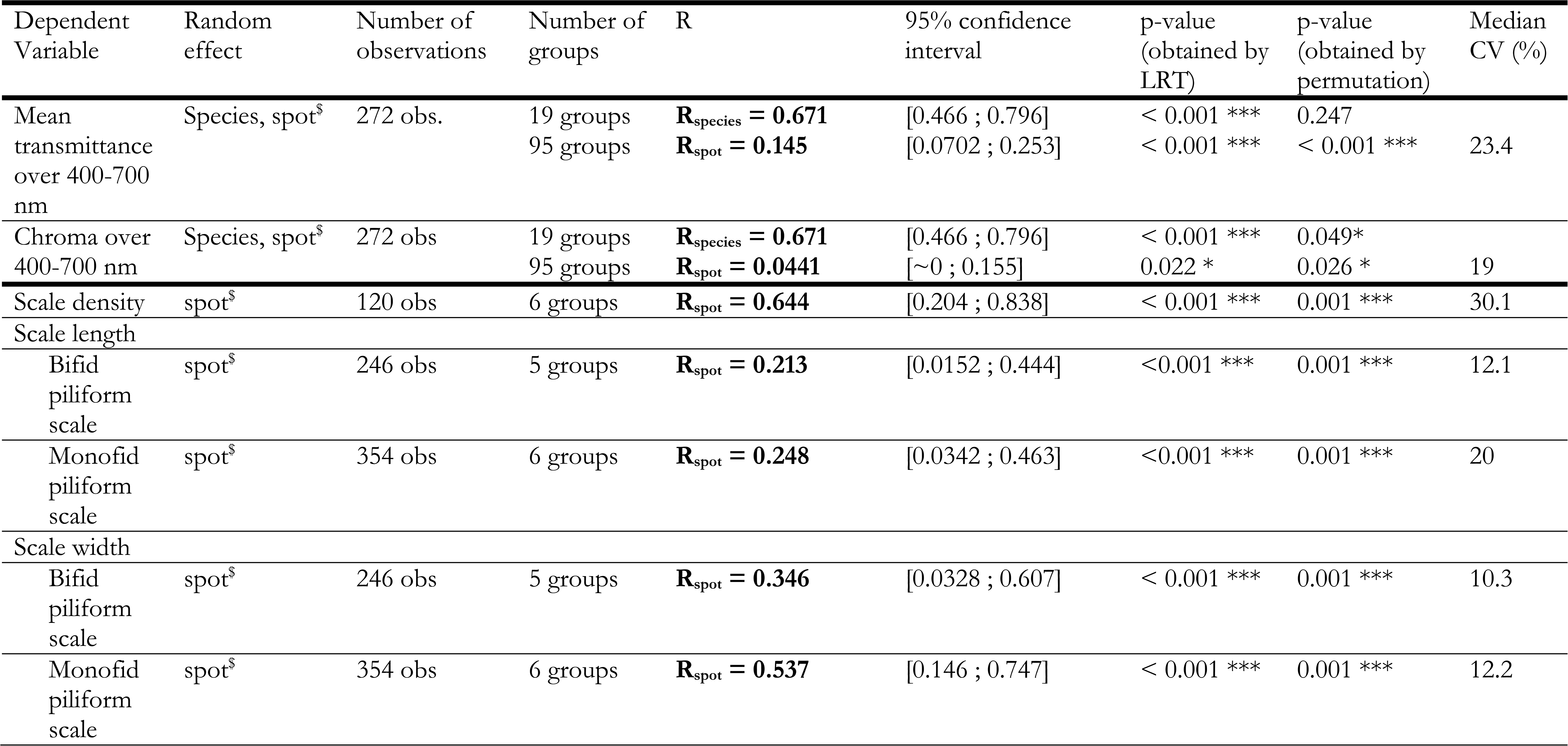
Repeatability of transmission measurements and structural features. For each grouping factor (either the number of species, or the number of different spots measured per species; indicated in the ‘number of groups’ column), we calculated the value of repeatability R based on several measurements of the same element of a grouping factor. The calculation of repeatability is based on mixed linear model. Confidence intervals are calculated with parametric bootstraping and p-values (associated to the test R > 0) are calculated with two methods : with likelihood ratio test comparing the likelihood of the model with and without the tested random effect and with permutation tests. We also calculated the coefficient of variation (CV, as the mean of the group devided by the standard error) for each group and we give here the median value of the CV distribution.

| Dependent Variable | Random effect | Number of observations | Number of groups | R | 95% confidence interval | p-value (obtained by LRT) | p-value (obtained by permutation) | Median CV (%) |
| --- | --- | --- | --- | --- | --- | --- | --- | --- |
| Mean transmittance over 400-700 nm | Species, spot<sup>s</sup> | 272 obs. | 19 groups<br>95 groups | $R_{\text{species}} = 0.671$<br>$R_{\text{spot}} = 0.145$ | [0.466 ; 0.796]<br>[0.0702 ; 0.253] | < 0.001 ***<br>< 0.001 *** | 0.247<br>< 0.001 *** | 23.4 |
| Chroma over 400-700 nm | Species, spot<sup>s</sup> | 272 obs | 19 groups<br>95 groups | $R_{\text{species}} = 0.671$<br>$R_{\text{spot}} = 0.0441$ | [0.466 ; 0.796]<br>[~0 ; 0.155] | < 0.001 ***<br>0.022 * | 0.049*<br>0.026 * | 19 |
| Scale density | spot<sup>s</sup> | 120 obs | 6 groups | $R_{\text{spot}} = 0.644$ | [0.204 ; 0.838] | < 0.001 *** | 0.001 *** | 30.1 |
| Scale length |  |  |  |  |  |  |  |  |
| Bifid piliform scale | spot<sup>s</sup> | 246 obs | 5 groups | $R_{\text{spot}} = 0.213$ | [0.0152 ; 0.444] | <0.001 *** | 0.001 *** | 12.1 |
| Monofid piliform scale | spot<sup>s</sup> | 354 obs | 6 groups | $R_{\text{spot}} = 0.248$ | [0.0342 ; 0.463] | <0.001 *** | 0.001 *** | 20 |
| Scale width |  |  |  |  |  |  |  |  |
| Bifid piliform scale | spot<sup>s</sup> | 246 obs | 5 groups | $R_{\text{spot}} = 0.346$ | [0.0328 ; 0.607] | < 0.001 *** | 0.001 *** | 10.3 |
| Monofid piliform scale | spot<sup>s</sup> | 354 obs | 6 groups | $R_{\text{spot}} = 0.537$ | [0.146 ; 0.747] | < 0.001 *** | 0.001 *** | 12.2 |

**Supplementary table 11.** Similarity between conspecific individuals for chromatic and achromatic contrasts. To test whether conspecific individuals were perceived as more similar than expected at random for each spot on the forewing, we randomized the contrasts over all pair of species and we calculated the mean distance for conspecific individuals. We compared the mean phenotypic distance (either chromatic or achromatic contrast) for the observed data to the distribution of mean phenotypic distance calculated for 10000 randomizations and we calculated the p-value as the number of randomizations where mean phenotypic distance was smaller than the observed phenotypic distance. We conclude that conspecific individuals are perceived as more similar than expected at random, implying that any individual is representative of its species.

| Dependant variable | Spot considered | factor | Observed mean contrast | p-value |
| --- | --- | --- | --- | --- |
| chromatic contrast dS | s1 | conspecific | 2.337709 | < 0.001 *** |
|  | s2 | conspecific | 2.700542 | < 0.001 *** |
|  | s3 | conspecific | 1.750813 | < 0.001 *** |
|  | s4 | conspecific | 1.831995 | < 0.001 *** |
|  | s5 | conspecific | 1.761623 | < 0.001 *** |
| achromatic contrast dL | s1 | conspecific | 3.236455 | < 0.001 *** |
|  | s2 | conspecific | 3.509987 | < 0.001 *** |
|  | s3 | conspecific | 1.838903 | < 0.001 *** |
|  | s4 | conspecific | 1.807439 | < 0.001 *** |
|  | s5 | conspecific | 1.651972 | < 0.001 *** |

**Supplementary result 1.**
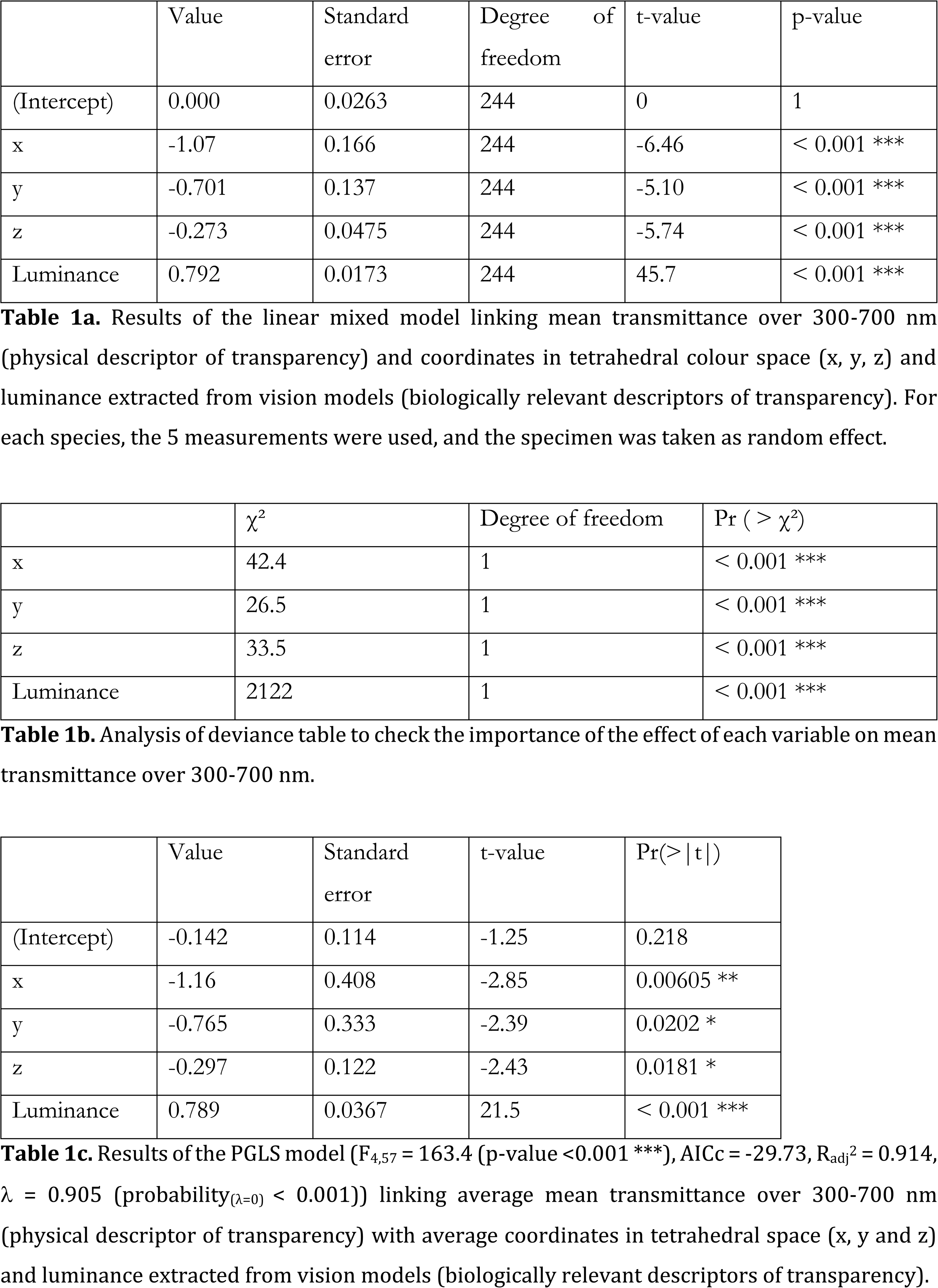
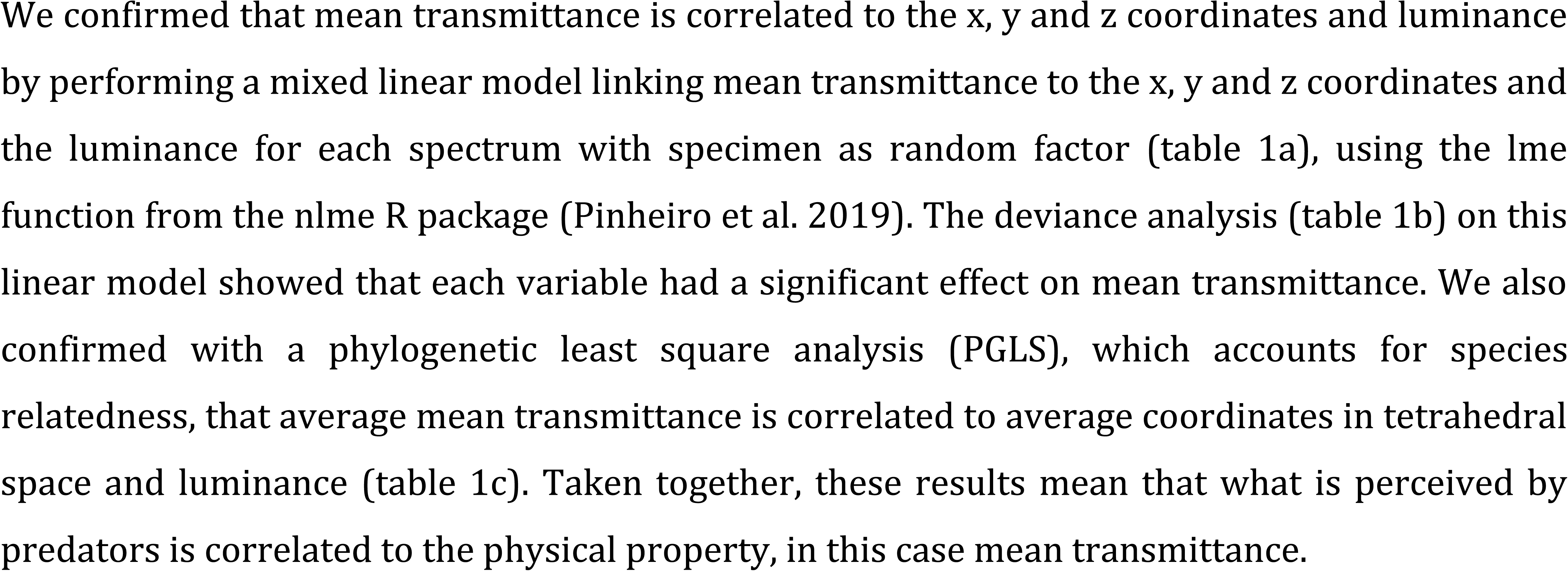
Link between physical and biologically relevant descriptors of transparency.

**Supplementary result 2.** Link between nanostructure density and nanostructure type.

|  | Sum of square | Mean square | F value | Pr (> F) |
| --- | --- | --- | --- | --- |
| Nanostructure type | 2341 | 585.3 | 26.26 | 0.001 |
| Residual | 1271 | 22.29 |  |  |

Results of the phylogenetic ANOVA on nanostructure density with nanostructure type as factor. The p-value is based on simulations.
|  | **Absence of nanostructures** | **Maze** | **Nipples** | **Sponge** | **pillar** |
| --- | --- | --- | --- | --- | --- |
| **Absence of nanostructures** | 0 (p =1) |  |  |  |  |
| **Maze** | 1.418 (p = 1) | 0 (p =1) |  |  |  |
| **Nipples** | 2.976 (p = 0.16) | 0.220 (p = 1) | 0 (p =1) |  |  |
| **Sponge** | **5.204 (p = 0.01)** | 2.167 (p = 0.62) | 3.800 (p = 0.89) | 0 (p =1) |  |
| **pillar** | **8.344 (p =0.01)** | **3.897 (p = 0.02)** | **8.588 (p = 0.01)** | **2.609 (p = 0.01)** | 0 (p =1) |

Results of the post-hoc tests (t values) to determine which type of nanostructure is different in density to others. P-value were corrected with Bonferroni correction and are indicated in brackets after t-values. Significant differences between nanostructure types are highlighted in bold.

## Supplementary Materials & Methods

### Phylogeny

We used published sequences from eight gene regions to infer a molecular phylogeny: the mitochondrial cytochrome oxidase c subunit 1 (COI) gene and the nuclear genes carbamyl-phosphate synthase II (CAD), malate dehydrogenase (MDH), elongation factor 1 alpha (EF-1α), tektin (TKT), ribosomal protein S5 (RpS5), isocitrate dehydrogenase (IDH) and Glyceraldehyde 3-phosphate dehydrogenase (GAPDH), which represent a total length of 7433 bp (Supplementary table 7). To improve phylogeny topology, we added 35 species representing 8 additional families to the dataset. When no sequence was available for a particular species on Genbank, we sequenced *de novo* the COI, CAD and MDH genes of that species (Supplementary table 7). We have missing data for some species, but we had at least the COI sequence for each species considered.

For *de novo* sequencing, DNA was extracted from butterfly legs with a DNeasy® Blood & Tissue Kit (QIAGEN laboratory) and targeted genes were amplified with PCR conditions adapted from Wahlberg and Wheat (2008). COI, CAD and MDH were amplified in two pieces with the primers described in Wahlberg and Wheat (2008). PCR were performed in a volume of 25 µL with 2 to 4 µL of genomic DNA, 1 µL of each primer at a concentration of 100 pmol/µL, 1 µL of nucleotides at a concentration of 2 mM, 2.5 µL of DreamTaq buffer, 0.125 µL of DreamTaq polymerase. The elongation phase was reduced to 70 seconds. For CAD and MDH, the annealing temperature was reduced to 50°C for most specimens. Eurofins Genomics sequenced the PCR products with Sanger method.

Sequences were aligned with CodonCodeAligner (version 3.7.1.1, CodonCode Corporation, http://www.codoncode.com/) and concatenated with PhyUtility (version 2.2, Smith and Dunn 2008). The dataset was then partitioned by gene and codon positions and the best models of substitution were selected over all models implemented in BEAST, using the ‘greedy’ algorithm and linked rates implemented in Partition Finder 1.0.1 (Lanfear et al. 2012, see Supplementary table 9 for best scheme). We performed a Bayesian inference of the phylogeny using BEAST 1.8.3 (Baele et al., n.d.) on the Cipres server (Miller et al., 2010). We constrained some clades to be monophyletic (notably Ithomiini, Danainae, Nymphalidae, Riodinidae, Pieridae, Papilionidae, Erebidae, Notodontidae, Geometridae, Noctuoidae, Papilionoidae) and we calibrated the crown age and divergence time of some groups (see Supplementary table 8), following Kawahara et al. (2019). Four independent analyses were run for 50 million generations, with one Monte Carlo Markov chain each and a sampling frequency of one out of 50 000 generations (resulting in 1000 posterior trees). After checking for convergence of the two best analyses, the posterior distributions of these two runs were combined (using logCombiner 1.8.2, Drummond and Rambaut 2007), with a burnin of 10%. The maximum clade credibility (MCC) tree with median node ages was computed using TreeAnnotator 1.8.2 (supplementary figure 5). Species not represented in our dataset were then pruned from the tree. The MCC tree was used for subsequent phylogenetic analyses.

### Spectrophotometry

Specular transmission was measured over 300-700 nm, a range of wavelength to which both butterflies and birds, which are supposed to be their main predators, are sensitive (Briscoe & Chittka, 2001; Hart, 2001) with a custom-built set-up composed of a 300W Xenon lamp emitting light over 200-1160 nm, a collimated emitting optic fibre (UV to NIR multimod fibre with a core diameter of 50 µm) illuminating the wing sample with a 1mm diameter spot and a collimated collecting optic fibre (Avantes UV to IR multimod fibre with a core diameter of 200 µm, FC-UVIR 200-1) connected to the spectrometer (SensLine AvaSpec-ULS2048XL-EVO, Avantes). Fibres are aligned and 14 cm apart. The wing is placed perpendicular to the fibres at equal distance with the ventral side facing the illuminating fibre. The spectrometer has a resolution of 0.5 nm and transmittance is calculated relative to a dark (light patch blocked at the end of the illuminating fibre) and to a white reference (no sample between the fibres):

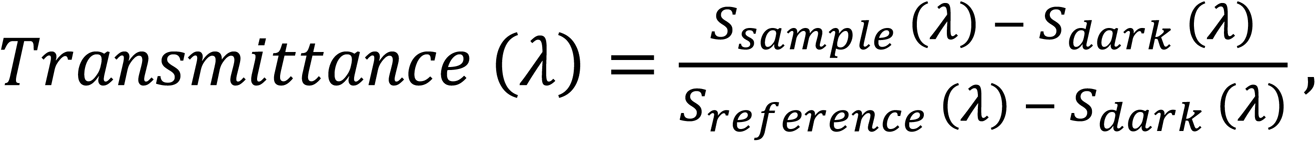

where λ represent the wavelength, S the number of photons counted by the spectrometer for this wavelength for sample, dark and reference measurements.

Each spectrum was smoothened with the loess function using R software (version 3.6.2.) (0.2 span on 500-700nm, and 0.05 on 300-700nm). We computed mean transmittance over 300-700 nm (B2) using Pavo2 (Maia et al., 2019), as a proxy for transparency: wing transparency increases as mean transmittance increases.

We assessed measurement repeatability on 11 species, representing part of six mimicry rings and belonging to 6 different families. We measured 3 times each of the five spots on the forewing (see supplementary figure 2 for location) for 2 to 3 individuals per species. We assessed measurement repeatability of mean transmittance (B2, Montgomerie 2006) and of chroma (S8, Montgomerie 2006) over 400-700 nm by calculating repeatability with species (11 groups for the 11 species considered), individual (32 groups for the 2 to 3 specimens per species) and spot (160 groups for the 5 spots measured for each of the 32 specimens) as random effect with rpt function from rptR package (Stoffel et al., 2017): R_species, B2_ = 0.586, p-value < 0.001; R_individual, B2_ = 0.0535, p-value = 0.033; R_spot, B2_ = 0.34, p-value < 0.001; R_species, S8_ = 0.583, p-value < 0.001; R_individual, S8_ = 0.0476, p-value = 0.053; R_spot, S8_ = 0.355, p-value < 0.001 (Supplementary table 10a).

We assessed intraspecific variation on 19 species, for which we had more than one individual, that represented six mimicry rings and belonged to 6 different families. We measured each of the five spots on the forewing once (see supplementary figure 2 for location) for 2 to 3 individuals per species. We assessed intraspecific variation of mean transmittance (B2, Montgomerie 2006) and of chroma (S8, Montgomerie 2006) over 400-700 nm by calculating repeatability with species (19 different groups) and spot (95 different groups corresponding to the 5 spot for each species) as random effect with rpt function from rptR package (Stoffel et al., 2017): R_species, B2_ = 0.671, p-value < 0.001; R_spot, B2_ = 0.145, p-value < 0.001; R_species, s8_ = 0.671, p-value < 0.001; R_spot, S8_ = 0.0441, p-value = 0.022 (Supplementary table 10b). As all measurements were repeatable, we considered that any individual is representative of its species. One specimen per species was therefore used in all analyses.

Due to technical issues, measurements for repeatability could only be performed for wavelengths ranging from 400 to 700 nm. However, both mean transmittance and chroma over 400-700 nm were highly correlated to mean transmittance and chroma over 300-700 nm, respectively (correlation coefficient equals 0.9979 [0.9974; 0.9983] (p-value < 0.001) for mean transmittance and 0.9765 [0.9707; 0.9812] (p-value < 0.001) for chroma). Repeatability in measurements over 400-700 nm can therefore be extrapolated to the full 300-700 nm range.

### High-resolution imaging and structure characterisation

Dry wings were cut from specimens before being gold-coated (10 nm thick layer) and observed in SEM (Zeiss Auriga 40). Top-view and cross section SEM images were analysed with ImageJ 1.52 (Schindelin et al., 2012) to measure scale density, scale length and width, membrane thickness, and nanostructure density. To measure scale density, we counted the number of scales in 3 different rectangular areas on a SEM photo and we calculated the mean scale density for each specimen. We measured scale length and width on 3 different scales, and we calculated the mean length and width for each specimen. We measured membrane thickness on 5 different photos per specimen and we calculated the mean membrane thickness. For nanostructure density, we coded macros (one for each type of nanostructure) in ImageJ (Schindelin et al., 2012) to count the number of structures on the image. For each specimen, we had two types of images with different magnifications. As the densities were congruent between the two types of images (measure of repeatability with rptR package: R = 0.368, p-value < 0.001), we calculated the mean nanostructure density between them.

We assessed intraspecific variation for scale characteristics (density, length and width) on three species, belonging to the mimicry ring agnosia: *Ithomia agnosia*, *Pseudoscada timna* and *Heterosais nephele.* We used 5 males and 5 females per species. We measured scale density for proximal and distal zone on forewing twice and we measured scale length and width for 10 different scales per specimen. We assessed repeatability with rptR with species as random effect (see Supplementary table 10): for density, R_species_ = 0.545 (p-value < 0.001); for scale length, R_species_ = 0.241 (p-value < 0.001); for scale width, R_species_ = 0.579 (p-value < 0.001). As scale structural features were repeatable within species, we used one specimen per species.

## Notes

### Competing Interest Statement

The authors have declared no competing interest.

